# The nematode *Caenorhabditis elegans* can benefit from its coexisting fungal microbiome member *Barnettozyma californica*

**DOI:** 10.1101/2025.02.17.638710

**Authors:** Carola Petersen, Hanne Griem-Krey, Christina Martínez Christophersen, Hinrich Schulenburg, Michael Habig

**Author notes:** These authors contributed equally (Carola Petersen, Hanne Griem-Krey). Corresponding author:* Michael Habig.

## Abstract

The nematode *Caenorhabditis elegans* is known to feed on and interact with bacteria in its environment and has become a model organism for microbiome studies. However, whether and how *C. elegans* interacts with co-occurring fungi remains largely unknown, despite the presence of many fungal species in its natural habitat. Here, we isolate the yeast *Barnettozyma californica* from a mesocosm experiment with *C. elegans* and characterize its genome and interaction with the nematode. We find that *B. californica* is ingested by *C. elegans* and can serve as a sole, albeit poor, food source. Yet, when present together with *Escherichia coli* OP50, the fungus can lead to higher population growth and altered foraging behavior, suggesting that this fungi-bacteria mixture provides a better food source than bacteria alone. This effect varied between different natural *C. elegans* strains, suggesting a genomic basis for the nematode’s interaction with *B. californica*. The fully assembled and annotated genome of the isolated *B. californica* strain does not indicate any obvious candidate genes for its interaction with *C. elegans* and/or *E. coli* OP50. Overall, our results provide an intriguing example of the complexity and multi-level relationship between naturally interacting fungi, bacteria, and animals.

**Graphical Abstract:** *Barnettozyma californica* is a member of the fungal microbiome of the model nematode *Caenorhabditis* elegans and can serve as food for worms. When combined with *E. coli*, this fungus can increase population growth and reduce foraging behaviour dependent on C. elegans genotype, revealing a genetic influence on host-fungus-bacterium interactions.

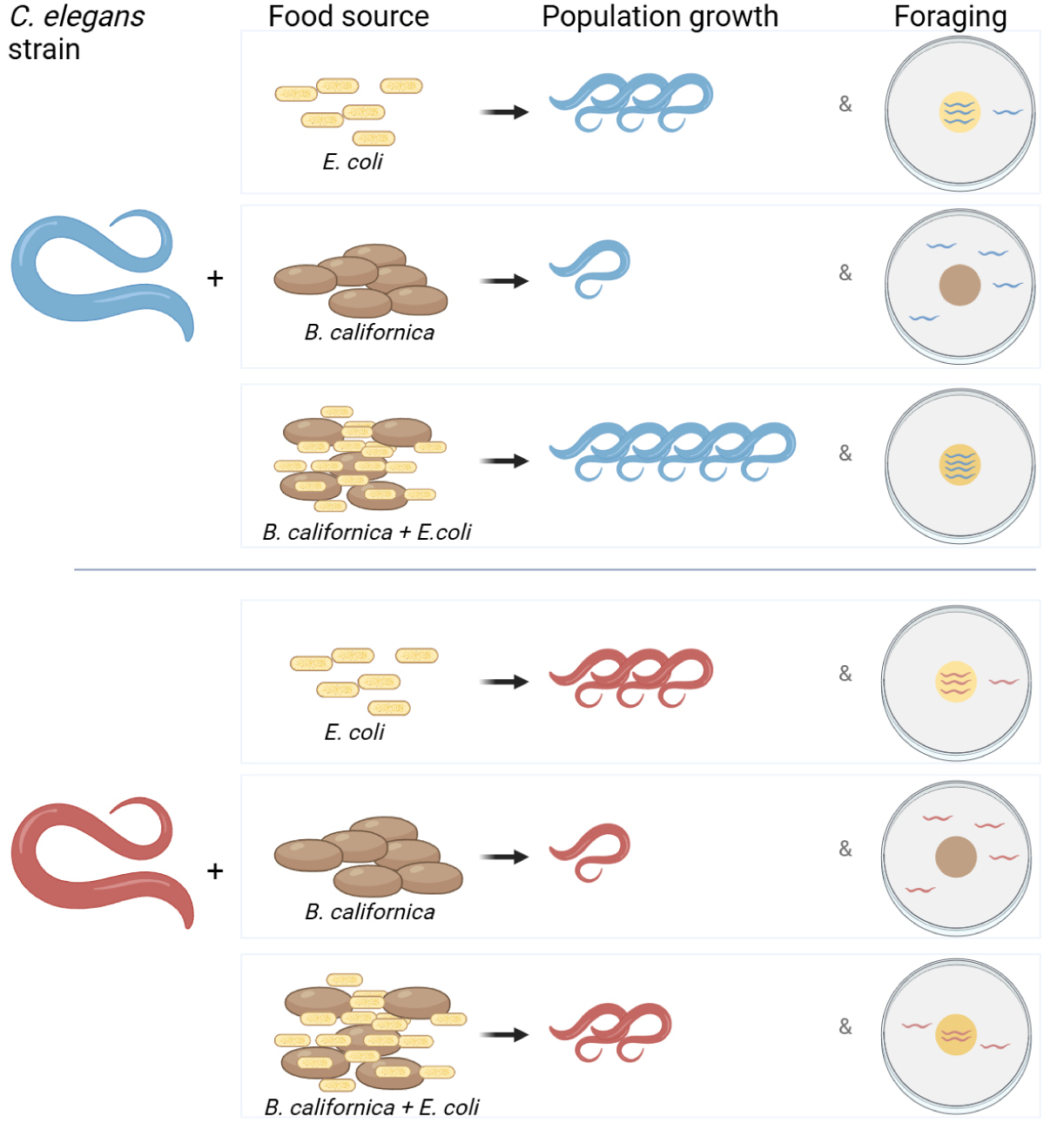

## Introduction

*Caenorhabditis elegans* inhabits a diverse microbial community in its natural environment, including bacteria and fungi. These microbes play various roles in the life cycle of *C. elegans* – serving as food, competitors, commensals, or pathogens – and can also be part of its microbiome. The interaction of *C. elegans* with pathogens such as the bacteria *Bacillus thuringiensis*, *Pseudomonas aeruginosa*, microsporidia of the genus *Nematocida*, the oomycete *Myzocytiopsis humicola*, or the Orsay virus have been studied over the last decades (Tan, Mahajan-Miklos and Ausubel, 1999; Troemel *et al*., 2008; Schulte *et al*., 2010; Félix *et al*., 2011; Osman *et al*., 2018; Tecle and Troemel, 2022; Tran and Luallen, 2024; Zárate-Potes, Schulenburg and Dierking, 2024). More recently, interest has expanded to include the worm’s bacterial microbiome with the first studies published in 2016 (Berg *et al*., 2016; Dirksen *et al*., 2016; Samuel *et al*., 2016; Schulenburg and Félix, 2017). These and subsequent studies have shown that the bacterial microbiome of *C. elegans* is distinct from its surrounding microbial environment under both natural and experimental conditions (Berg *et al*., 2016; Dirksen *et al*., 2016, 2020; Samuel *et al*., 2016; Petersen *et al*., 2023; Zimmermann *et al*., 2024; Johnke *et al*., 2025). In *C. elegans*, unclassified Enterobacteriaceae are the most common bacteria, along with members of the genera *Pseudomonas*, *Stenotrophomonas*, *Ochrobactrum*, and *Sphingomonas* (Zhang *et al*., 2017). The composition, assembly, stability, and variation of the microbiome are influenced by the host’s developmental stage and genotype (Dirksen *et al*., 2016; F. Zhang *et al*., 2021; Zimmermann *et al*., 2024). The majority of bacterial strains isolated from either the worm’s habitat or directly from its intestine support robust *C. elegans* growth. However, some can be pathogenic (Samuel *et al*., 2016; Griem-Krey *et al*., 2023; Gonzalez and Irazoqui, 2024). To better understand the influence of microbial communities on the host, a simplified synthetic 12-member microbiota community, known as the CeMbio resource, was developed (Dirksen *et al*., 2020). All CeMbio bacteria can colonize the *C. elegans* intestine, either as a community or individually, to varying degrees. Most strains accelerate the worm’s development compared to those grown on standard lab food, *Escherichia coli* OP50 (Dirksen *et al*., 2020). However, some CeMbio bacteria shorten the lifespan of immunodeficient animals (Gonzalez and Irazoqui, 2024). Beneficial effects, such as the protective role of *Pseudomonas lurida* MYb11 against the pathogen *B. thuringiensis* Bt679, can be disrupted by p38 MAPK signaling inhibition. This suggests that microbiota effects on *C. elegans* can be context-dependent (Kissoyan *et al*., 2019, 2022; Griem-Krey *et al*., 2023; Pees *et al*., 2024).

Like bacteria, many fungi are found in the natural habitat of *C. elegans* (Dirksen *et al*., 2016). In contrast to its interactions with naturally co-occurring bacteria, very little is known about its interactions with co-occurring fungi. The focus has primarily been on pathogenic interactions. Spores of *Drechmeria coniospora* and other natural fungal endoparasites of *C. elegans*, such as *Hirsutella rhossiliensis*, *Haptoglossa dickii*, and *Catenaria anguillulae*, infect nematodes and complete their vegetative life cycle within infected hosts. Upon infection, *D. coniospora* produces enterotoxins in the epidermis of *C. elegans* which manipulate the host’s immune responses (X. Zhang *et al*., 2021). Additionally, *C. elegans* has been used as a model organism to study hosts immune responses towards fungal human pathogens (Madende *et al*., 2020). Innate immune responses of *C. elegans* towards fungal pathogens involve antimicrobial peptides, lectins, lysozymes, the p38 and ERK mitogen-activated protein kinase pathways, as well as the DAF-2 and TGF-ß/DBL-1 signaling pathways (Zugasti and Ewbank, 2009; Zugasti *et al*., 2016). Moreover, *C. elegans* produces chemical compounds with antifungal activity, making it a valuable infection model (Breger *et al*., 2007; Madende *et al*., 2020). A completely different type of interaction between fungi and nematodes involves nematode-trapping fungi, carnivorous microorganisms that capture and digest nematodes using specialized trapping structures (Jiang, Xiang and Liu, 2017). These include *Arthrobotrys* spp. and *Drechslerella stenobrocha*, which ensnare nematodes with constricting ring traps or adhesive traps (Yang *et al*., 2007; Nordbring-Hertz, Jansson and Tunlid, 2011; Liu *et al*., 2012), highlighting the diverse range of possible interactions between fungi and nematodes.

Apart from these few well-characterized interactions between fungi and nematodes, and despite the fact that fungi have been isolated directly from natural *C. elegans* strains (Dirksen *et al*., 2016), little is known about fungi as members of the nematode’s microbiome. In a recent study examining *C. elegans* in laboratory compost mesocosms, several fungal species were identified, with *Barnettozyma californica* strains being particularly abundant in the worms (Petersen *et al*., 2023). Interestingly, worm populations that lived for multiple generations in a microbial community that included *Barnettozyma* showed a particularly high fitness. This raises the question of whether the observed fitness advantage was directly caused by *Barnettozyma* fungi and whether this fungal taxon plays an important role for *C. elegans* – either as a food source or as a member of its microbiota.

*Barnettozyma* is a genus in the order Phaffomycetales, which was established in 2023 based on a phylogenetic analysis of 290 BUSCO genes (Groenewald *et al*., 2023). The genus *Barnettozyma* was originally described in 2008 based on a multigene phylogenetic analysis, which included *B. californica* (previously known as *Williopsis californica*) (Kurtzman, Robnett and Basehoar-Powers, 2008). *B. californica* has a worldwide distribution (Kurtzman, 2011), with its type strain CBS 252 first isolated from soil in California in 1931 (Lodder, 1932). Although relatively few genomic sequences are available for *B. californica*, the first draft genomes – produced using short-read sequencing technologies – were obtained from two *B. californica* isolates from Ireland (Mullen *et al*., 2018). To date, no chromosome-level genome assemblies exist for this species. *B. californica* has demonstrated the ability to grow on xylose and can utilize a broader range of carbon sources than other tested members of the *Barnettozyma* genus. It also exhibits ammonium reduction capabilities and can utilize both nitrite and nitrate (Falih and Wainwright, 1995; Kurtzman, 2011; Shen *et al*., 2018). Additionally, *B. californica* strain K1, isolated from surface sediments of the Pearl River, has been shown to perform heterotrophic nitrification-aerobic denitrification (Fang et al., 2021). Recently, another member of the *Barnettozyma* genus, *B. botsteinii*, was discovered in the intestinal tract of the termite *Macrotermes bellicosus*. Like *B. californica*, *B. botsteinii* can grow on xylose, and its ability to degrade plant-derived polymers suggests a potential role in assisting termite metabolism (Arrey *et al*., 2021). In summary, despite its wide distribution and presence in various environments, many aspects of *B. californica*’s life history and genomic characteristics remain to be explored. Additionally, how *B. californica* interacts with *C. elegans* and whether and how it directly affects the nematode’s fitness is unknown.

Here, we characterize the interaction between *B. californica* and *C. elegans* and show that *B. californica* can serve as a food source and influence the behavior of *C. elegans*. When provided as the sole food source, it is less beneficial than bacterial food but can still support the complete life cycle of *C. elegans*. In contrast, when *B. californica* is offered as food in combination with bacteria – presumably more representative of *C. elegans*’ natural environment – it can increase *C. elegans* fitness, though this effect depends on the nematode’s genotype. Thus, we demonstrate that a fungus co-occurring with *C. elegans* can influence the worm’s fitness.

## Results

We first isolated *B. californica* MYf642 from a mesocosm experiment and characterized its interaction with *C. elegans*. We then analyzed the genome of *B. californica* MYf642 to determine whether it reflects its interaction with *C. elegans*.

### *C. elegans* ingests *B. californica* MYf642

Worms exposed to the microbiome of a specific compost mesocosm consistently showed a higher relative abundance of ASVs from the fungal genus *Barnettozyma* (Petersen *et al*., 2023). We isolated *B. californica* MYf642 from this mesocosm and tested whether *C. elegans* ingests it. To visualize *B. californica* MYf642, we used calcofluor white staining, which specifically binds to chitin. We also used the *C. elegans* strain dkIs37[*act-5p::GFP::pgp-1*], in which the intestinal basal membrane is fluorescently labeled to aid localization. Our results show that *C. elegans* ingests *B. californica* MYf642, with fungal cells observed throughout the whole intestine, posterior to the grinder (Fig. 1, Fig. S1). However, it is not entirely clear whether intact *B. californica* cells can pass through the grinder unaltered. Additionally, we observed and documented feeding behavior of *C. elegans* including ingestion, passage through the worm intestine and excretion of *B. californica* MYf642 (Movies S1-2). Control worms fed solely on *E. coli* OP50 did not show any stained cells in their intestine because *E. coli* does not contain chitin (Fig. S1). Additionally, previous studies have shown that intact *E. coli* cells are not found posterior to the grinder in *C. elegans*.

**Figure 1:**
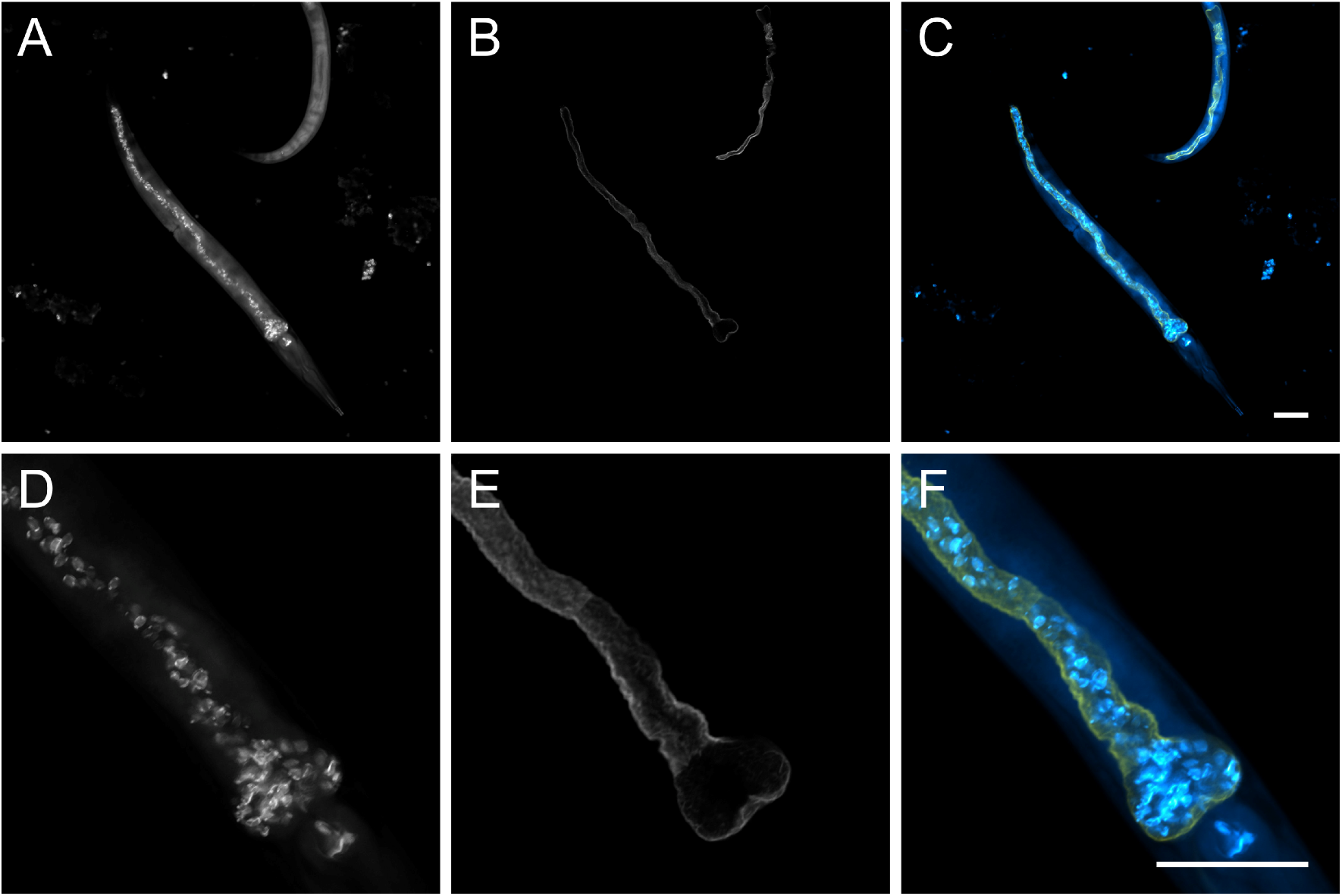
*C. elegans* ingest *Barnettozyma californica* MYf642. (A–F) *C. elegans* dkIs37[*act-5p::GFP::pgp-1*] can ingest *B. californica* MYf642, with fungal cells observed throughout the intestine (A, C, D, F) (white or light blue). The apical intestinal membrane of *C. elegans* dkIs37[*act-5p::GFP::pgp-1*] expresses GFP and is shown in white (B, E) or yellow (C, F). Calcofluor white (CFW) stained *B. californica* MYf642 is depicted in white (A, D) or light blue (C, F). Worms were visualized using confocal laser scanning microscopy. Scale bars represent 50 µm. Images were false-colored using ImageJ.

### *B. californica* MYf642 is a poor food source for *C. elegans* alone, but can significantly increase the fitness when presented alongside bacterial food

Next, we tested whether *B. californica* can serve as a sole food source or, in combination with the standard laboratory food source *E. coli* OP50, support the entire life cycle and influence the fitness of *C. elegans*. To account for the genetic variability of *C. elegans* and avoid restricting our analysis to the canonical laboratory strain N2, we included eleven additional strains from the *C. elegans* mapping population provided by the *Caenorhabditis* Natural Diversity Resource (CaeNDR) (Cook *et al*., 2017; Crombie *et al*., 2024). In a population growth assay, three L4 larvae of each *C. elegans* strain were added to microbial lawns consisting of either *B. californica* MYf642, *E. coli* OP50, or a combination of both (which would, by containing both a fungal and a bacterial component, presumably be more akin to the microbial composition in the natural habitat of *C. elegans*). The number of offspring was determined after five days (Fig. 2A). For all *C. elegans* strains tested, *B. californica* MYf642 as the sole food source resulted in significantly lower population growth compared to the *E. coli* OP50 control. On average, the number of offspring on *B. californica* ranged from 0.4% to 5.1% of the respective offspring count on *E. coli* OP50, with a maximum of an average of 79 offspring per original worm in strain CB4856 and a minimum of 20 on average in strain JU775. Although more than three adult worms were observed after five days – indicating that *B. californica* MYf642 can support the complete life cycle of *C. elegans* – the low number of offspring per worm compared to on *E. coli* suggests that *B. californica* is a poor food source on its own.

**Figure 2:**
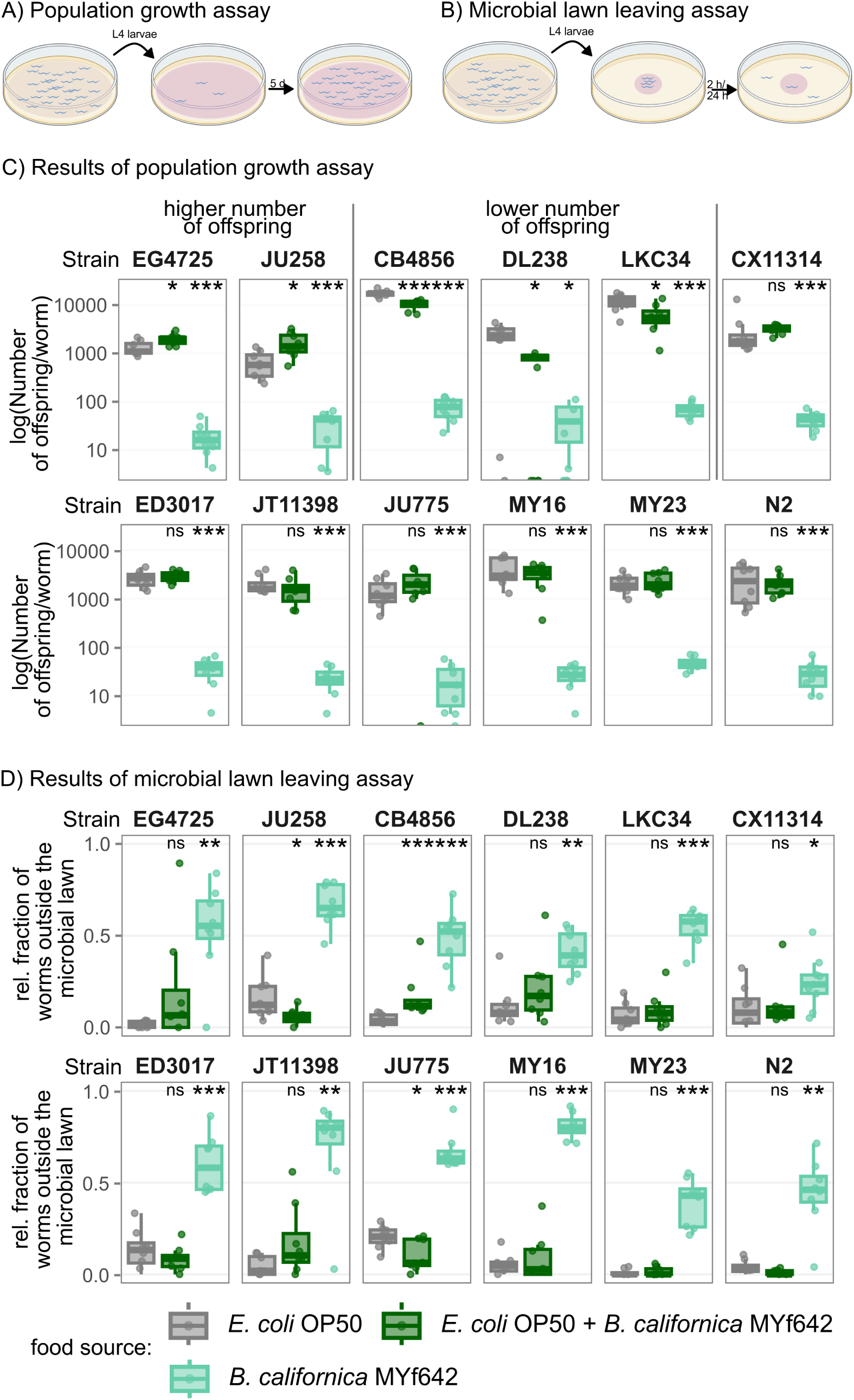
*B. californica* MYf642 in combination with *E. coli* OP50 as food results in a worm strain-dependent effect on the number of *C. elegans* offspring and behavioral response. A) Outline of the population growth assay: Three synchronized L4 larvae are transferred onto a microbial lawn and the population size is evaluated after 5 days. B) Outline of the microbial lawn leaving assay: Approximately 30 synchronized L4 larvae are put onto a central microbial lawn and after 2 h and 24 h the fraction of nematodes outside the microbial lawn is determined. C-D) The figure shows the effect on population growth (C) and the behavioral response (D) on *E. coli* OP50 (gray), *E. coli* OP50 combined with *B. californica* MYf642 (dark green), and *B. californica* MYf642 alone (light green). *B. californica* MYf642 as the sole food source results in significantly lower number of offspring per worm and a higher number of worms outside the microbial lawn after 24 h across all worm strains (compared to *E. coli* OP50 as sole food source). When combined with *E. coli* OP50, *B. californica* MYf642 leads to a higher number of offspring in *C. elegans* strains EG4725 and JU258, while significantly lowering the number of offspring in strains CB4856, DL238, and LKC34. In JU258 and CB4856, this is associated with fewer or more worms leaving the microbial lawn, respectively. Significant differences (determined by the Wilcoxon rank sum test with Holm correction for multiple testing) compared to *E. coli* OP50 as the sole food source are indicated as follows: *p* < 0.05 (*), *p* < 0.01 (**), *p* < 0.001 (***), ns = non-significant. n = 8

The poor population growth of *C. elegans* on *B. californica* MYf642 alone contrasts with the results observed when the fungus was combined with bacterial food (Fig. 2A). On the combined microbial lawn, for eight *C. elegans* strains, population growth was not significantly different from that on the *E. coli* OP50 control. For three other strains (CB4856, DL238, and LKC34), the number of offspring was significantly lower on the mixed microbial lawn, reaching 59%, 19%, and 53% of the respective *E. coli* OP50 control levels. Most interestingly, for *C. elegans* strains EG4725 and JU258, the number of offspring was significantly higher on the mixture of *E. coli* OP50 and *B. californica* MYf642. Here, the mean number of offspring per worm was 150% and 252% of the respective *E. coli* OP50 control, indicating a strong positive effect on population growth. Using population growth as a proxy for fitness, we conclude that *B. californica* MYf642 can influence the fitness of *C. elegans* when present alongside a bacterial food source. This effect varies from positive to negative, depending on the *C. elegans* genotype.

### *B. californica* MYf642 affects the behavior of *C. elegans* in a genotype-dependent manner

Next, we hypothesized that *B. californica* might influence the exploratory behavior of the nematode, similar to how different bacterial species affect dietary choice behavior in *C. elegans* (Shtonda and Avery, 2006; Meisel and Kim, 2014). To test this, we placed approximately 30 synchronized L4 worms onto a microbial lawn consisting of either *B. californica* MYf642, *E. coli* OP50, or a mix of these two. The microbial lawn-leaving behavior of twelve *C. elegans* strains was evaluated by measuring the fraction of worms found outside the microbial lawn after 2 h and 24 h (Fig. 2B for 24 h data and Fig. S2 for 2 h data). When placed on a *B. californica* MYf642 lawn, a significantly larger fraction of worms was found outside the microbial lawn after 24 h compared to on the *E. coli* OP50 control. This suggests that the nematodes increase their foraging behavior due to *B. californica* being a poor food source. This contrasts with the situation when the microbial lawn contained both the fungi and the bacteria. Here, we observed *C. elegans* genotype-dependent variation in the response. After 24 h, a significantly higher fraction of worms left the mixed microbial lawn in strain CB4856 – one of the strains that also exhibited lower population growth on these mixed microbial lawns. Interestingly, a lower fraction of worms left the mixed microbial lawn for the *C. elegans* strains JU258 (which displayed higher population growth on these mixed microbial lawns) and JU775. This suggests that, for these *C. elegans* strains, the mixed microbial lawn may provide a more favorable environment. Although there appears to be a correlation between the results of the microbial lawn-leaving assay at 24 h and the population growth assay (Fig. S3), this correlation is not statistically significant. In addition to the 24-h time point, we also tested lawn-leaving behavior after 2 h. Interestingly, at this earlier time point, four *C. elegans* strains – including CB4856, which at 24 h showed a much larger fraction outside the mixed microbial lawn – remained more on the mixed microbial lawn than on the bacterial lawn. In summary, *B. californica* MYf642 affects the foraging behavior of *C. elegans* when mixed with *E. coli* OP50 in a genotype-dependent manner.

### The genome of *B. californica* MYf642 comprises seven chromosomes and one mitochondrial genome

Having established that *B. californica* MYf642 affects the fitness and behavior of *C. elegans*, we next sequenced and assembled its genome as a resource for future experimental analyses and also to identify patterns that might help us understand its interaction with *C. elegans*. Using PacBio HiFi CLR reads we compared three different assembly methods (see Methods and Materials). The best assembly contained seven nuclear contigs and one mitochondrial genome, resulting in a total genome size of 12.1 Mb (Fig. 3A, Table S1). A full genomic phylogeny based on orthologous proteins identifies the isolate MYf642 to fall within the species *B. californica* (Fig. S4A). We compared the eight nuclear contigs with four publicly available, more fragmented assemblies of *B. californica* and found a high degree of synteny between orthologous genes (Fig. 3B, Fig. S4B, C). With one exception, no contig from the fragmented assemblies spans more than one contig of the *B. californica* MYf642 assembly. The exception is the fragmented assembly of *B. californica* strain PB4207, where one contig shows synteny with both TIG09 and TIG15 of the assembly. However, this synteny discrepancy is also present compared to the other three fragmented assemblies of *B. californica*, suggesting that it is either due to a structural variation or an assembly error in PB4207. This conclusion is supported by the presence of telomeric repeats at both ends of five of the seven contigs assembled here by us (Fig. 3A). Although we did not detect the canonical 5’-TTAGGG-3’ telomeric repeat, we identified a novel telomeric repeat sequence, 5’-GTGTGG-3’, which differs from the wide variety of telomeric repeats previously described in ascomycetous yeasts, especially species within the subphylum *Saccharomycotina* (Červenák *et al*., 2021). The telomeric repeat sequence we identified is similar to but shorter than those described for other members of the *Phaffomycetaceae* (Červenák *et al*., 2021). The presence of telomeric repeats at both ends of five contigs, along with the strong synteny across all published *B. californica* genomes, supports our conclusion that *B. californica* MYf642 has seven nuclear chromosomes.

**Figure 3:**
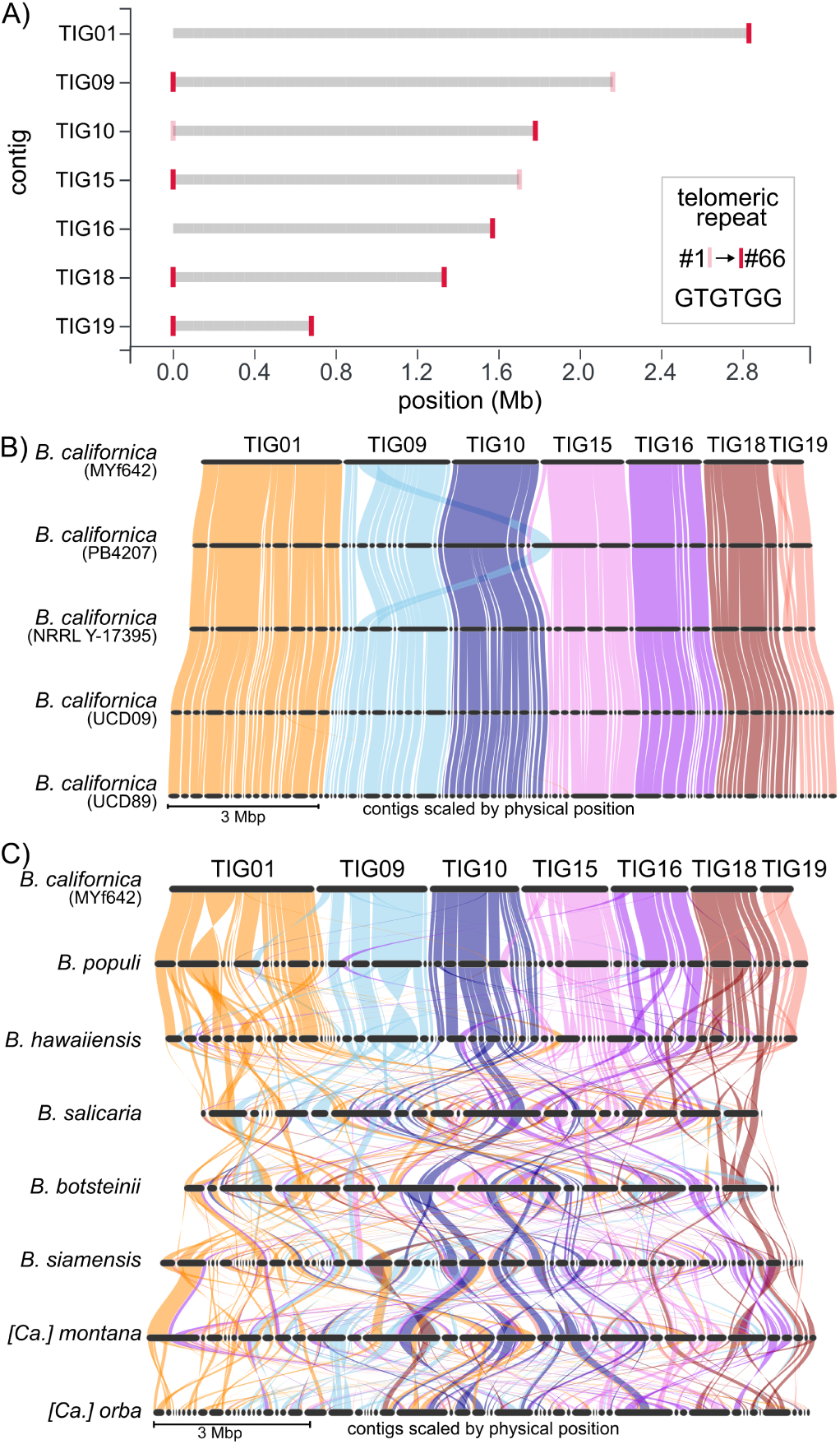
Chromosome-level assembly of *B. californica* MYf642. A) Tapestry report on contig sizes and the presence of telomere repeats in the seven contigs of the *B. californica* MYf642 assembly. Five of the seven contigs show telomeres at both ends. B) Synteny plot based on orthologous genes between *B. californica* MYf642 and four publicly available fragmented assemblies of *B. californica*, showing high levels of synteny across all *B. californica* MYf642 contigs. The absence of syntenic regions spanning more than one contig or scaffold suggests that the *B. californica* MYf642 contigs represent individual chromosomes (see also Fig. S7). C) Synteny plot between *B. californica* MYf642 and other species within the *Barnettozyma* genus reveals high structural variation, which increases with phylogenetic distance.

### The genome of *B. californica* MYf642 is densely populated with genes and contains a low proportion of repetitive and transposable elements

We annotated TEs, genes, tRNAs, and functionally characterized putative secreted proteins, effectors, proteases, and Carbohydrate-Active Enzymes (CAZymes), along with gene-wise relative synonymous codon usage (Tables S1 and S2). The genome of *B. californica* MYf642 contains 5,893 genes, comparable to the 5,802–5,803 genes reported in the fragmented assemblies of *B. californica* strains UCD09 and UCD5803 (Mullen *et al*., 2018). The genome is gene-dense, with genes covering 70.5% of the genome, while TEs account for a relatively low 2.1%. Of the gene transcripts, 312 contain a secretion signal, and among these, a relatively small number (78) are putative effectors – small secreted proteins that are thought to interact with potential hosts (Jones and Dangl, 2006) or other microorganisms (Snelders *et al*., 2020). Additionally, we identified 191 putative proteases, 143 putative CAZymes, and a total of 428 TEs and repeats (Table S2). Among the TEs, *hAT* transposons, a widespread class II DNA transposon (Atkinson, 2015), are the most frequent in the *B. californica* MYf642 genome (Table S3). Furthermore, we functionally annotated all transcripts using Blast2GO (Table S4).

### The genome of *Barnettozyma californica* MYf642 is similar to other members of the *Barnettozyma* genus

We next compared the synteny of genes from our chromosome-level assembly with high-continuity genome assemblies of other *Barnettozyma* species to assess whether the high synteny observed within *B. californica* also extends to other species in the genus. Our results show that this is not the case. Instead, we find a low level of synteny between different *Barnettozyma* species. The closely related *B. populi* shows a relatively high level of synteny, with few inversions and translocations. In contrast, more distantly related species show much lower levels of synteny (Fig. 3C and Fig. S4B). We, therefore, conclude that synteny across species within the *Barnettozyma* genus is generally low. We also compared the genome functional composition of 74 additional species within the *Phaffomycetales* order, including *Komagataella pastoris* as an outgroup to discern patterns of evolution and adaptation in the family. Notably, *Komagataella* was previously classified within the *Phaffomycetaceae* (Kurtzman, 2011), but genome-scale analyses now place it under the *Pichiales*, making it a suitable outgroup for this study (Shen *et al*., 2018; Groenewald *et al*., 2023). To correlate phylogenetic relationships with genome functional composition, we reconstructed a maximum likelihood phylogenetic tree based on concatenated protein sequences of single-copy orthologous genes (a total of 657 genes) (Fig. 4, right). To ensure comparability between the species, we functionally reannotated each of the assemblies using the same bioinformatics pipeline employed for the annotation of *B. californica* MYf642. Our results confirm the monophyletic origins of the genera within the *Phaffomycetaceae*, specifically *Barnettozyma*, *Cyberlindnera*, *Phaffomyces*, and *Starmera*. However, the *Wickerhamomycetaceae*, which contains the genus *Wickerhamomyces*, does not appear monophyletic and is interspersed among species of the *Phaffomycetaceae*. Although outside the main focus of this study, it is important to note that the non-monophyletic characteristics of the *Wickerhamomyces* genus, which we here inferred from 657 orthologous genes, contrast with earlier analyses based on internal transcribed spacer (ITS) and large ribosomal subunit (LSU) sequences (Arrey *et al*., 2021; Nundaeng *et al*., 2021), or elongation factor-1α (EF-1α) and small ribosomal subunit (SSU) sequences (Kobayashi, Kanti and Kawasaki, 2017). The functional genome composition of *B. californica* MYf642 is similar to that of other species within the genus *Barnettozyma* and other species in the *Phaffomycetales*. Despite a high variation in certain genomic features, such as transposable element content (ranging from 0.2% to 7.4%), secreted proteins (239–482), effector candidates (58–248), proteases (175–238), and CAZymes (112–172), none of these variations appear to correlate with phylogenetic relationships, nor does *B. californica* MYf642 stand out as an extreme in any category. Additionally, we observed high variation in codon usage bias, as predicted by gene-wise relative synonymous codon usage (RSCU), and the predicted numbers of tRNAs. However, neither codon usage bias nor tRNA numbers appeared to be correlated or follow any phylogenetic pattern, and again nor does *B. californica* MYf642 stand out as an extreme. In conclusion, the functional genome composition of *B. californica* MYf642 is broadly similar to other related species, despite the observed variations in certain genomic traits.

**Figure 4:**
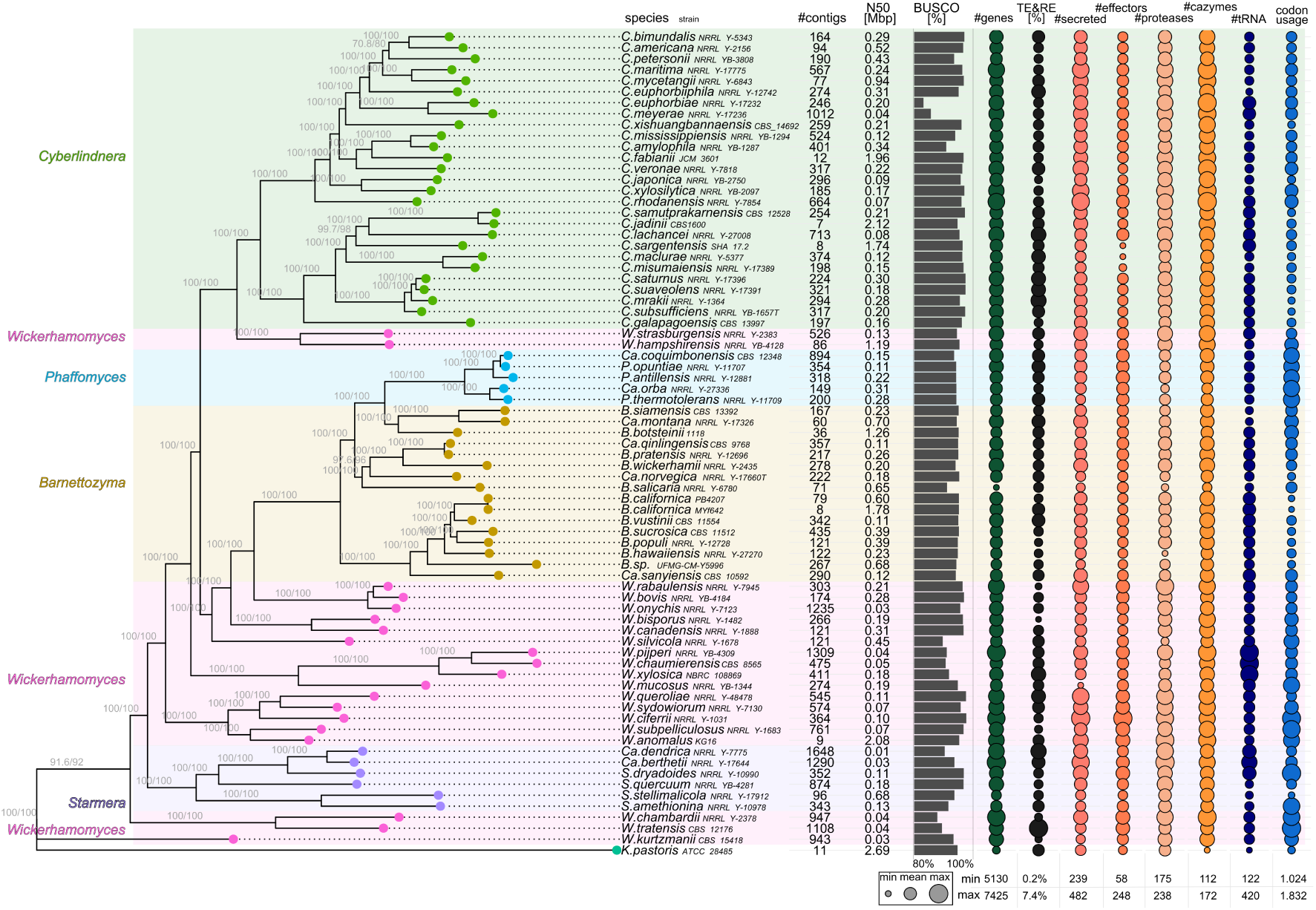
Phylogeny and summary of functional annotation of Phaffomycetales. Left: Maximum Likelihood (ML) phylogenetic tree of single-copy orthologous proteins from the genera *Starmera*, *Phaffomyces*, *Cyberlindnera*, *Barnettozyma*, and *Wickerhamomyces*, with *Komagataella pastoris* used as an outgroup. The genus *Wickerhamomyces* (pink), part of the family *Wickerhamomycetaceae*, is polyphyletic and interspersed among other genera from the family *Phaffomycetaceae*. Right: High variation is observed in the presence of transposable elements (TEs), secreted proteins, putative effectors, putative CAZymes, and proteases. The number of TEs and codon usage patterns vary greatly between species and do not correlate with phylogenetic relationships.

### *B. californica* MYf642 contains unique genes associated with terpenoid metabolism and transport

We next compared the gene composition of *B. californica* MYf642 with representative members of other genera in the *Phaffomycetales* for which high-continuity genome assemblies are available. A total of 4,258 orthogroups were shared across all six species, while 28 orthogroups were unique to *B. californica* MYf642 (Fig. 5A, B). Based on these 28 orthogroups and functional gene annotations (Table S4), we conducted a GO-term enrichment analysis. This analysis identified several significantly over-represented GO terms, including “terpenoid metabolic processes,” “positive regulation of (R)-carnitine transmembrane transport,” and “polyamine transmembrane transport.” Interestingly, terpenoids and terpenes, form the largest group of secondary metabolites in fungi (González-Hernández *et al*., 2023). Additionally, we tested whether genes involved in nitrification and denitrification, previously reported in the *B. californica* K1 strain – specifically amoA, nirK, nosZ (Fang *et al*., 2021) – were present in *B. californica* MYf642. However, none of these genes were detected by BLAST searches using the sequences described in (Fang *et al*., 2021). Therefore, we conclude that *B. californica* MYf642 does not possess the genetic capacity for heterotrophic nitrification-aerobic denitrification.

**Figure 5:**
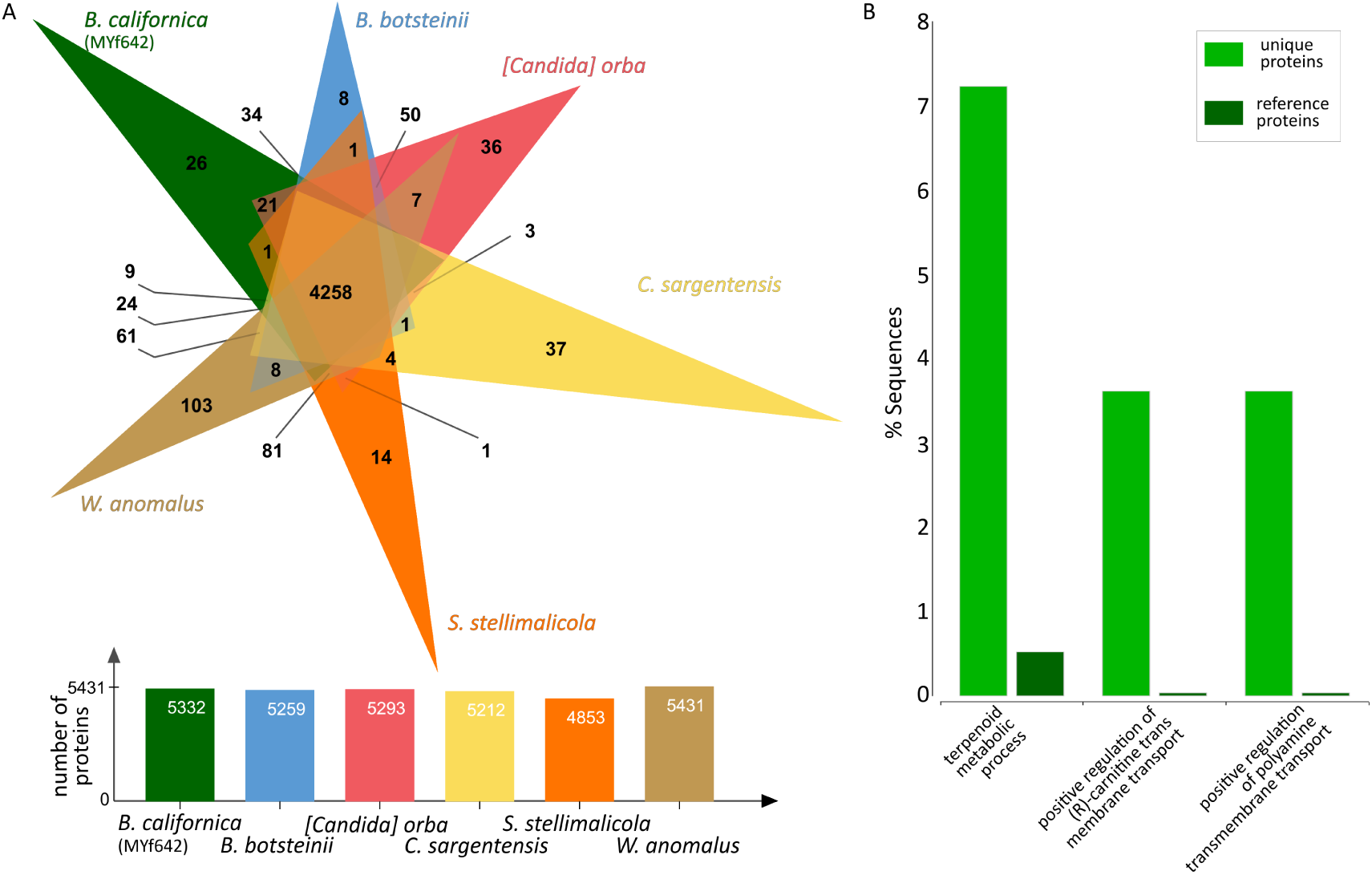
Identification of proteins restricted to *B. californica* MYf642 and associated GO terms. A) Venn diagram of orthologous proteins among representative species of the Phaffomycetales compared to the *B. californica* MYf642 isolate, identifying 28 clusters with a total of 83 proteins that are restricted to *B. californica* MYf642 and do not have orthologs in the other representative species. B) Results of the GO term enrichment analysis for the 83 proteins restricted to *B. californica* MYf642 compared to the reference set (all proteins encoded by the *B. californica* MYf642 genome), with significance thresholds of *p* = 0.05 and FDR = 0.1.

## Discussion

Here, we show that the fungus *B. californica*, which is found in diverse environments and was isolated as a putatively beneficial microbiome member of *C. elegans* from a mesocosm experiment, is ingested and used as food by *C. elegans*. On its own, it appears to be a poor food source, leading to increased *C. elegans* foraging behavior. However, when combined with a bacterial food source, it can have a positive fitness effect and reduce foraging activity. Furthermore, we report the chromosome-level assembly of the *B. californica* MYf642 genome, which includes a novel type of telomeric repeat. The genome is highly compact and does not exhibit any distinct features that set it apart from other *Phaffomycetales* yeasts and that could thus explain its interaction with *C. elegans* and its bacterial food.

The interaction between *C. elegans* and fungi is poorly understood, and the extent to which fungi contribute to the *C. elegans* diet remains unknown. Our study is one of the few that demonstrate that fungi can serve as a food source for *C. elegans*. Similarly, the basidiomycete yeasts *Cryptococcus kuetzingii* and *C. laurentii* have been used as *C. elegans* food sources in the lab (Mylonakis *et al*., 2002). However, for neither *Cryptococcus* species was a direct association with *C. elegans* shown, nor were they isolated from a *C. elegans* environment. Notably, in our experiments, the positive effect on *C. elegans* population growth was observed only when *B. californica* MYf642 was present together with *E. coli* OP50. This suggests that, in addition to directly affecting *C. elegans* biology, *B. californica* MYf642 may influence it indirectly via interactions with *E. coli* OP50. In general, fungi and bacteria that share habitats can interact through the production of antibiotics, signalling molecules, and other compounds, such as siderophores that manipulate nutrient availability (Deveau *et al*., 2018; Scherlach and Hertweck, 2020; Pierce *et al*., 2021). Currently, the mechanism underlying the positive effects of *B. californica* MYf642 in the presence of *E. coli* OP50 remains unclear. One possibility is that the fungus produces compounds that are limited in a pure bacterial culture. Many fungi are known to produce secondary metabolites with great structural diversity (Macheleidt *et al*., 2016), which may be beneficial to *C. elegans*. Alternatively, and not mutually exclusively, *B. californica* may induce changes in *E. coli* that are beneficial to *C. elegans*. In any case, based on our results, this effect also appears to be dependent on the genotype of the nematode.

We also found that the foraging behavior of *C. elegans* was affected by *B. californica* MYf642 when present in combination with *E. coli* OP50, and this effect was again dependent on the *C. elegans* genotype. It is known that genetically different *C. elegans* strains vary in their lawn-leaving behavior in response to food bacteria and pathogens (Reddy *et al*., 2009; Bendesky *et al*., 2011; Nakad *et al*., 2016). Leaving a microbial lawn can be an avoidance behavior toward certain pathogens, such as *Serratia marcescens* (Pradel *et al*., 2007) or *B. thuringiensis* (Schulenburg and Müller, 2004; Hasshoff *et al*., 2007; Nakad *et al*., 2016). However, *C. elegans* can also distinguish between high- and low-quality food bacteria. Worms tend to search for high-quality food that supports growth, while they avoid or abandon low-quality food (Shtonda and Avery, 2006). Thus, the increased foraging behavior in the presence of *B. californica* MYf642 indicates that it is a poor food source, while the reduced foraging behavior of some *C. elegans* strains when present with the mixture of fungi and bacteria indicates that this is a good food source. In microbial lawn-leaving assays, nematodes often return to the food multiple times within minutes to hours, which might explain the differences between our 2-h and 24-h data in the lawn-leaving assay.

The genomes of fungi are shaped by interactions with other organisms that share the same habitat, including other microorganisms. The discovery of penicillin by Alexander Fleming in 1929 was the first indication that fungi manipulate their microbial environment through what we now recognize as secondary metabolites. Fungi also produce small secreted proteins, so-called effectors. Until recently, the role of effectors in plant-associated fungi was largely thought to be limited to interactions with the host and the suppression of the host immune system (Jones and Dangl, 2006). More recently, effectors have been shown to influence the bacterial and fungal microbiome of plants, suggesting that this may have been their ancestral function (Snelders *et al*., 2018, 2020, 2021). Although we were unable to identify any unique secondary metabolite clusters or effectors in the genome of *B. californica* MYf642 that might explain the fungus’s positive impact – both in this study and in the mesocosm study (Petersen *et al*., 2023) – this does not rule out the existence of yet-undiscovered genetic components involved in these interactions. Additionally, it is puzzling that we did not find substantial genetic overlap with the termite-associated *B. botsteinii* (Arrey *et al*., 2021). Thus, the fungal genetic components underlying the interaction with *C. elegans* and *E. coli* OP50 remain unknown.

In conclusion, both the microbial lawn-leaving assay and the population growth assay demonstrated that *B. californica* MYf642 affects *C. elegans* behavior when presented jointly with bacteria, in a genotype-dependent manner. The use of genetically diverse mapping strains from the CaeNDR has allowed us to identify these patterns, which are not apparent in the standard laboratory N2 strain. The interaction between *C. elegans*, *B. californica* MYf642, and *E. coli* OP50 that we present here is an intriguing example of the complexity and multi-level nature of interactions between fungi, bacteria, and animals – for which *C. elegans* is one of the ideal model organisms.

## Supplementary Figures

**Figure S1:**
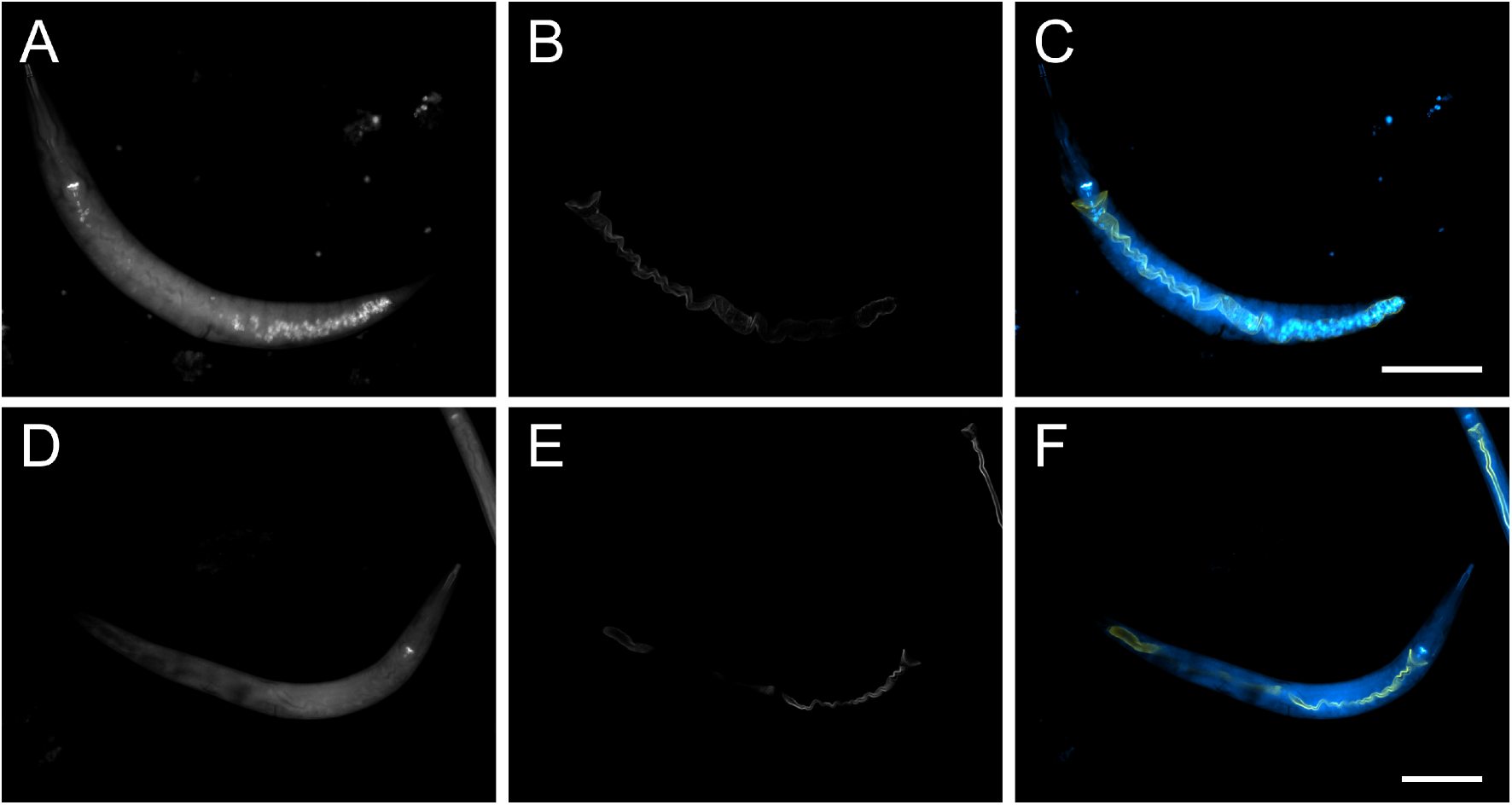
Stained *Barnettozyma californica* MYf642 but no stained *E. coli* OP50 in the *C. elegans* intestine. *B. californica* MYf642 can be found in various regions of the *C. elegans* intestine (A, C) (white or light blue) while *E. coli* OP50 cells do not contain chitin and hardly pass the worm grinder and are therefore not stained by the calcofluor white staining (D, F). The apical intestinal membrane of *C. elegans* dkIs37[*act-5p::GFP::pgp-1*] expresses GFP and is shown in white (B, E) or yellow (C, F). Calcofluor white (CFW) stained *B. californica* MYf642 is shown in white (A, D) or light blue (C, F). Worms were visualized using confocal laser scanning microscopy. The scale bars have a length of 100 µm. Images were false-colored using ImageJ.

**Figure S2:**
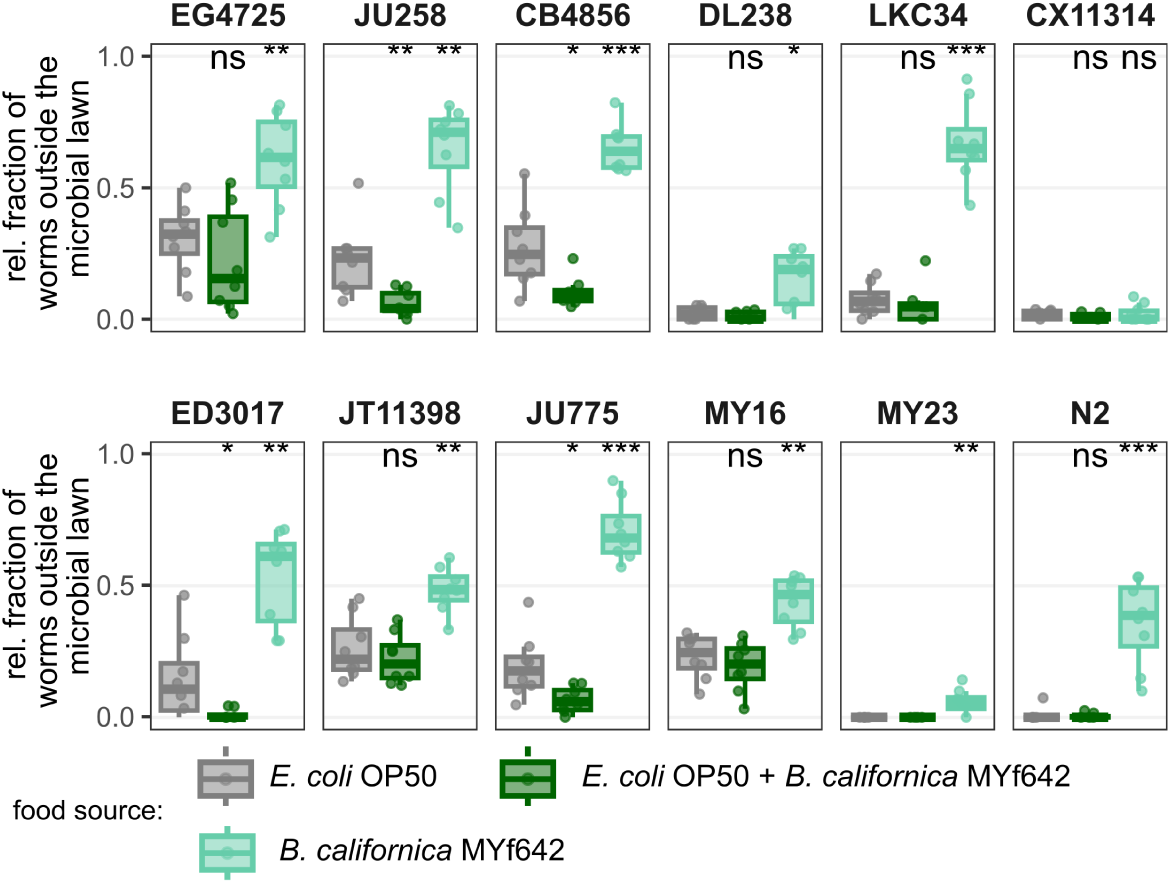
*B. californica* MYf642 in combination with *E. coli OP50* as food results in a strain-dependent effect on the behavioral response after 2 h. The figure shows the behavioral response of the indicated *C. elegans* strains on *E. coli* OP50 (gray), *E. coli* OP50 combined with *B. californica* MYf642 (dark green), and *B. californica* MYf642 alone (light green). *B. californica* MYf642 as the sole food source results in a higher number of worms outside the microbial lawn after 2 h for all strains (compared to *E. coli* OP50 as sole food source), with the exception of CX11314. When combined with *E. coli* OP50, *B. californica* MYf642 leads to genotype dependent variation in the fraction of worms outside the microbial lawn. Significant differences (determined by the Wilcoxon rank sum test with Holm correction for multiple testing) compared to *E. coli* OP50 as the sole food source are indicated as follows: *p* < 0.05 (*), *p* < 0.01 (**), *p* < 0.001 (***), ns = non-significant. n = 8

**Figure S3:**
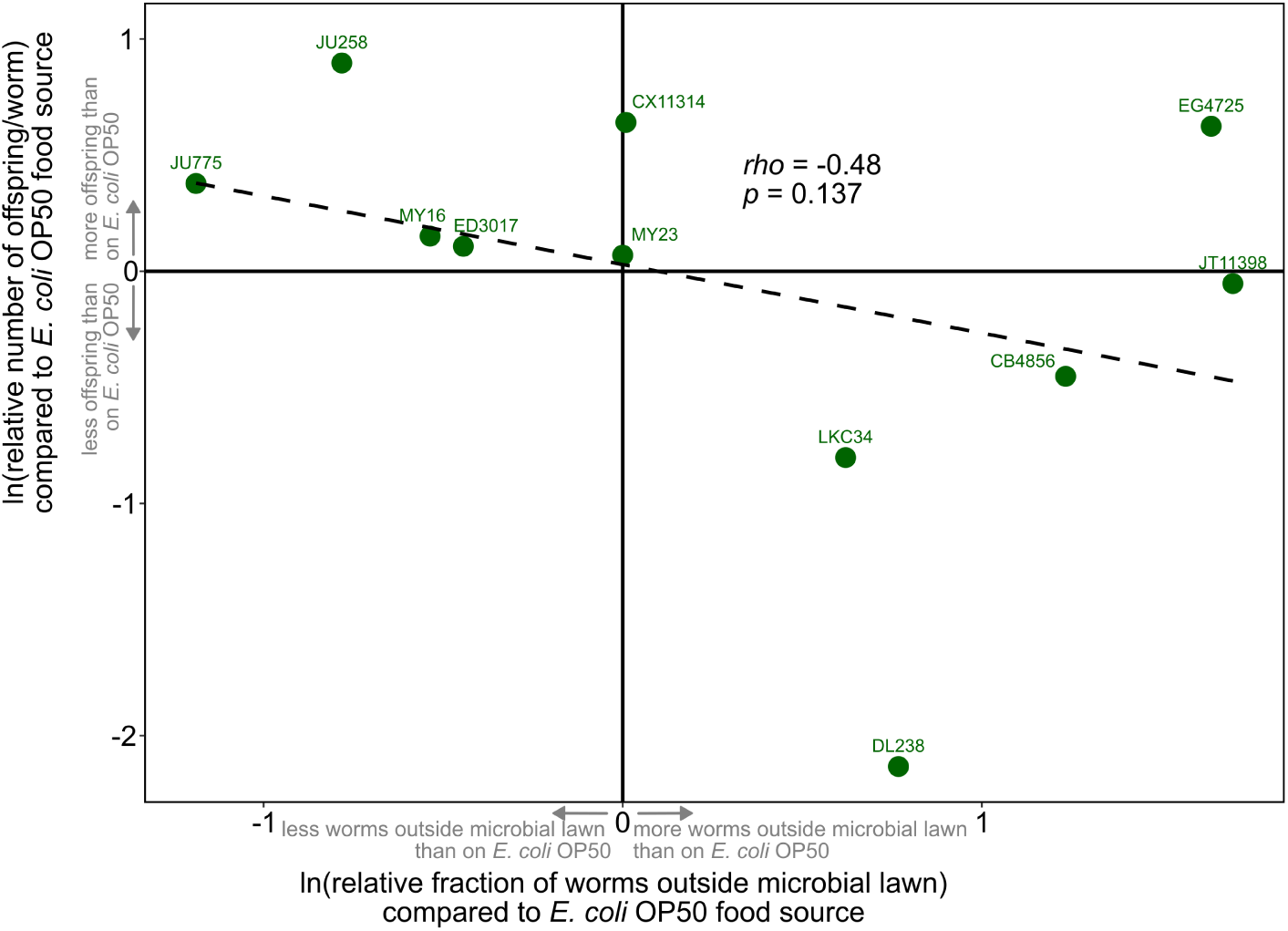
Correlation between lawn-leaving behavior and population growth rate. The relative number of offspring per worm for each *C. elegans* strain on the mixed microbial lawn (normalized to the *E. coli* OP50 control) is plotted against the relative fraction of worms outside the microbial lawn for each *C. elegans* strain (normalized to the *E. coli* OP50 control). A linear regression line is indicated by the dashed line.

**Figure S4:**
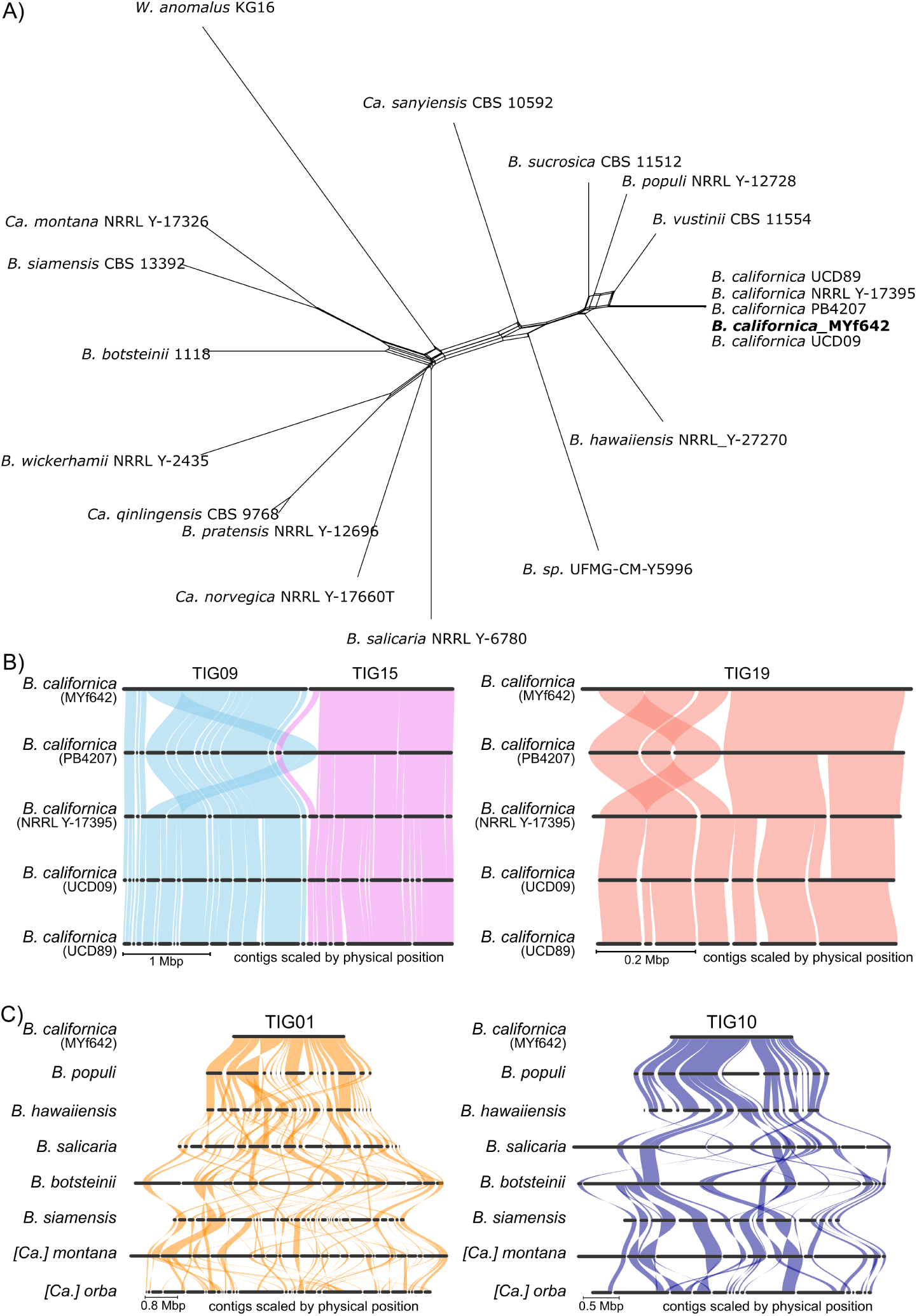
Network phylogeny of species within the genus *Barnettozyma* and details on synteny within *B. californica* support chromosome-level assemblies and extensive structural variation within the *Barnettozyma* genus. A) NeigborNet phylogeny of 2710 single copy orthologous proteins of species within the genus *Barnettozyma* and with *E. anomalus* as an outgroup. The isolate *B. californica* MYf642 falls within the species *B. californica*. B) Details of orthologous proteins based synteny for TIG09 and TIG15 (left) and TIG19 (right) show that the apparent structural variation is unique to the *B. californica* strain PB4207, where it is located at the end of a contig. All other previously published assemblies show no structural variation compared to *B. californica* MYf642. Hence this apparent structural variation in *B. californica* strain PB4207 could be due to a misassembly. C) Details of synteny for TIG01 (left) and TIG10 (right) demonstrate extensive structural variation, which increases with phylogenetic distance. Panels B) and C) show details from Figure 3 panel B) and C), respectively.

## Supplementary Tables

Table S1: Overview of *B. californica* MYf642 genome assembly and characteristics

Table S2: Comparison of the functional annotations of the genomes used for the phylogenetic analysis

Table S3: Overview of the transposable elements (TE) and repeat elements (RE) in the B. californica MYf642 assembly

Table S4: Functional annotation of proteins encoded by *B. californica* MYf642 Table S5: Genome assembly accessions included in this study

## Supplementary Movies Captions

Movie S1: Feeding behavior of *C. elegans* including ingestion, passage through the worm intestine and excretion of *B. californica* MYf642

Movie S2: Feeding behavior of *C. elegans* including ingestion, passage through the worm intestine and excretion of *B. californica* MYf642

## Methods and Materials

### Microbial and *C. elegans* strains

*E. coli* OP50 and *B. californica* MYf642 were cultured overnight in a shaking incubator (180 rpm) in LB medium at 37 °C or 28 °C, respectively. We used the 12 *C. elegans* strains CB4856, CX11314, DL238, LKC34, ED3017, EG4725, JT11398, JU258, JU775, MY16, MY23, and N2 from a mapping population provided by the *Caenorhabditis* Natural Diversity Resource (CaeNDR) (Cook *et al*., 2017; Crombie *et al*., 2024). Additionally, the *C. elegans* strain dkIs37[*act-5p::GFP::pgp-1*] was used to locate stained cells of *B. californica* MYf642 in the *C. elegans* intestine. All *C. elegans* strains were maintained on nematode growth medium (NGM) on *E. coli* OP50.

### Isolation of *Barnettozyma californica* MYf642

We isolated *B. californica* MYf642 from a 100-day-old laboratory compost mesocosm, originating from a long-term evolution experiment with *C. elegans* and its associated microbes. Worms from this 100-day-old laboratory compost showed improved fitness and enrichment of *Barnettozyma* in a follow-up common garden experiment when exposed to the microbial community from day 100 (Petersen *et al*., 2023). To isolated *Barnettozyma californica* MYf642, a random compost sample was mixed with 25 ml M9 with 0.025% Triton X-100 (M9-T). A 300 µl aliquot of this mixture was homogenized with 10 zirconia beads (1 mm) in a Bead Ruptor 96 (OMNI International, Kennesaw, GA, USA) at 30 Hz for 3 min, then frozen in 10% dimethyl sulfoxide (DMSO) at −80 °C. Thawed material was serially diluted (10⁻¹ to 10⁻⁴) in sterile phosphate buffered saline (PBS), and 50 µl of each dilution was plated onto potato dextrose agar (39 g/l, Carl Roth, Karlsruhe, Germany). Plates were incubated at room temperature for 10 days. Morphologically distinct colonies were picked upon appearance and purified by re-culturing for at least three times. Pure isolates were grown in lysogeny broth (LB) on a circular shaker for 2 days at 25 °C and preserved in 35% glycerol at −80 °C. Crude DNA was extracted by adding 5 µl liquid culture to 19.5 µl 5x PCR buffer with 0.5 µl proteinase K (20 mg/ml), followed by freezing at −80 °C for 30 min, digestion at 50 °C for 1 h and heat inactivation at 95 °C for 15 min. For identifying yeasts and fungi, 1 µl crude DNA served as template in a 50 µl PCR reaction using DreamTaq DNA polymerase (Thermo Fisher Scientific, Darmstadt, Germany) to amplify the internal transcribed spacer (ITS) region with the primers ITS1f (5’-CTTGGTCATTTAGAGGAAGTAA-3’) and ITS4r (5’-TCCTCCGCTTATTGATATGC-3’) (White *et al*., 1990; Gardes and Bruns, 1993).

A one-day culture of the *B. californica* MYf642 isolate in LB medium, grown at 28 °C and 200 rpm, was used for DNA isolation following a modified version of the cetyltrimethylammonium bromide (CTAB) DNA extraction protocol (Plissonneau, Stürchler and Croll, 2016). Briefly, the cells were harvested by centrifugation, washed twice, and 400 µl of the cell pellet was crushed using a mortar and pestle with liquid nitrogen. Following extraction with CTAB, DNA was isolated twice using phenol:chloroform:isoamylalcohol and precipitated with isopropanol. The DNA was washed three times with ethanol before being resuspended in 1x TE buffer. Sequencing was performed using the PacBio Revio platform at BGI Techsolutions (Hong Kong), resulting in 308,037 CCS reads with a mean length of 17,689 bp.

### Calcofluor white staining

Calcofluor white stain is a fluorochrome that binds to cellulose and chitin in cell walls of fungi. L4 larvae of the *C. elegans* strain *dkIs37[act-5*p::GFP*::pgp-1]* expressing PGP-1::GFP in the apical intestinal membrane (Sato *et al*., 2007) were transferred to nematode growth medium (NGM) plates inoculated with *B. californica* MYf642. After 24 h, adult worms were washed from the plates with M9-buffer containing 0.3% Tween20 (M9-T20). 500 µl of the worm-containing M9-T20 was added to a 2-ml tube, and 500 µl of 10 mM tetramisole was added to stop the ingestion and excretion of microbes. After about 2 minutes, the pellet of immobilized worms was transferred to a new tube. The worms were fixed and stained by adding 250 µl of calcofluor white staining solution consisting of equal parts calcofluor white stain and ethanol. After 3 h, the staining solution was removed, the worms were washed three to four times with M9-T20 and then examined for stained *B. californica* MYf642 in their intestine using confocal laser scanning microscopy (ZEISS LSM 880).

### Population growth assay

The population growth rate is a measure for worm fitness as it combines information on the developmental time, reproductive rate and survival. To start the assay, three *C. elegans* larvae in the fourth larval stage (L4) were picked to a 1-ml lawn of either *B. californica* MYf642, *E. coli* OP50 or a 1:1 mixture of both at OD_600_ 10 on 9 cm peptone-free medium plates. After 5 days at 20 °C, the worms were washed from the plates to 15 ml Falcon tubes using 5 ml M9-T. The liquid in the Falcon tubes was weighed for later calculations and then frozen at −20 °C until the worms were counted. Three subsamples were counted per sample, from which a mean value was calculated to account for variations in worm numbers. The total number of offspring over five days was calculated by dividing the mean worm count of the three subsamples by the number of counted microliters multiplied by the total weight of the sample in gram. The offspring number was divided by the three original worms to obtain the offspring per worm.

### Microbial lawn leaving assay

*C. elegans* can perceive the quality of its food as well as the presence of pathogens and adapt its behavior accordingly (Shtonda and Avery, 2006; Meisel and Kim, 2014). To determine the behavior of *C. elegans* towards *B. californica* MYf642, we performed a microbial lawn leaving assay. Approximately 30 synchronized L4 worms were pipetted to a 60 µl microbial spot of *B. californica* MYf642, *E. coli* OP50 or a 1:1 mixture of both adjusted to OD_600_ 10 on 6 cm peptone-free medium plates. The number of worms on and outside the spot was counted after 2 h and 24 h to analyze the early and late lawn leaving behavior. The escape rate was then calculated by dividing the number of worms outside the microbial spot by the total number of counted worms.

### Bioinformatics Analysis

#### Genome assembly

The code used for all subsequent bioinformatics analyses has been deposited here: https://github.com/michaelH-git/BarnettozymaCalifornica_genomeAnalysis.git. Briefly, assemblies were generated using Canu (2.2) (Koren *et al*., 2017), HiFiasm (0.19.9-r616) (Cheng et al., 2021), and Flye (2.9.3-b1797) (Kolmogorov et al., 2019). The assembly quality was checked using QUAST (v5.2.0) (Mikheenko *et al*., 2018). After visual inspection, the Canu-based assembly was selected for all subsequent analyses. The quality of this assembly was further evaluated using Tapestry (1.0.1) (Davey *et al*., 2020), and contigs that showed no coverage by the original reads were discarded (40 contigs, totaling 1.3 Mb). Of the remaining nine contigs, the smallest one (TIG20) showed synteny to mitochondrial genomes. This sequence was further polished and annotated using MitoHiFi (3.2.1) (Uliano-Silva *et al*., 2023), resulting in a 48,643 bp circular contig representing the mitogenome. For the remaining eight contigs representing the nuclear genome, telomere sequences were identified using TelFinder (Q *et al*., 2023), and the quality of the genome was rechecked using Tapestry.

#### Functional annotation of genomes

The list and accessions of the genome assemblies included in this study is given in Table S5. Genomes were functionally annotated using the following tools. Transposable and repetitive elements were identified using EDTA (2.2.0) (Ou *et al*., 2019). In cases where no transposable elements (TEs) were detected, repetitive sequences were called using EarlGrey (4.4.4) (Baril, Galbraith and Hayward, 2024). These annotations were used to softmask the genomes for gene annotation. Genes were annotated using BRAKER3 (3.0.8) (L *et al*., 2024) with protein evidence from OrthoDB11 restricted to fungi. Secondary metabolite clusters were annotated using antiSMASH (7.1.0) (Blin *et al*., 2021), and functional annotation of predicted proteins was performed using eggNOG-mapper (2.1.12) (Cantalapiedra *et al*., 2021) and InterProScan (Blum *et al*., 2021). Genome completeness was assessed with BUSCO (5.7.1) (Simão *et al*., 2015), while tRNAs were predicted using tRNAscan-SE (2.0.12) (Chan *et al*., 2021). Codon usage was estimated by determining the gene-wise relative synonymous codon usage (gRSCU) with BioKit (Steenwyk and Buida, 2022). Putatively secreted proteins were identified using SignalP (5.0b) (Almagro Armenteros *et al*., 2019), and putative effectors were predicted using EffectorP3 (3.0) (Sperschneider and Dodds, 2022). Putative CAZymes (carbohydrate-active enzymes) were identified using dbCAN (Zhang *et al*., 2018). All these annotations were combined using Funannotate (*Funannotate*, 2017). For *B. californica* MYf642 the functional annotation was further improved using Blast2GO (Conesa and Götz, 2008). Fisher’s Exact test for GO-term enrichment was conducted using this improved annotation in Blast2GO.

#### Phylogenetic analysis

Identification of single copy orthologs for all genomes of the phylogenetic analysis was conducted using Orthofinder (Emms and Kelly, 2019) restricted to one transcript for each gene. All single copy orthologous proteins shared among all species (657) were concatenated and aligned using MAFFT and the phylogenetic tree determined by IQTree using the LG+I+G model and generating a consensus tree of 1000 bootstrapings (Minh *et al*., 2020). Determination of orthologous proteins between *B. californica* MYf642, *B. botsteinii* (GCA_020280145.1), *C. sargentensis* (GCA_020995425.1), *S. stellimalicola* (GCA_030580055.1), *W. anomalus* (GCA_019321675.1) and *[Candida] orba* (GCA_003708145.3) to represent all genera within the phylogenetic analysis was performed using OrthoVenn3 (Sun *et al*., 2023). GO term enrichment analysis was performed on those genes unique to *B. californica* MYf642. Synteny analysis was conducted using GENESPACE (v1.3.1) (Lovell *et al*., 2022).

## Data availability

Sequencing reads, genome assemblies and annotations for *B. californica* MYf642 are deposited using Bioproject PRJNA1224554.

## Acknowledgements

CP and HS are thankful for support by the CRC1182 projects A1.1 and A4.3 funded by the German Science Foundation. The graphical abstract and Fig2 were created in BioRender. Habig, M. (2025) https://BioRender.com/y15j225 https://BioRender.com/d81s260

## References

Almagro Armenteros, J.J. et al. (2019) ‘SignalP 5.0 improves signal peptide predictions using deep neural networks’, Nature Biotechnology, 37(4), pp. 420–423. Available at: 10.1038/s41587-019-0036-z.

Arrey, G. et al. (2021) ‘Isolation, characterization, and genome assembly of Barnettozyma botsteinii sp. nov. and novel strains of Kurtzmaniella quercitrusa isolated from the intestinal tract of the termite Macrotermes bellicosus’, G3: Genes|Genomes|Genetics, 11(12), p. jkab342. Available at: 10.1093/g3journal/jkab342.

Atkinson, P.W. (2015) ‘hAT Transposable Elements’, in *Mobile DNA III*. John Wiley & Sons, Ltd, pp. 773–800. Available at: 10.1128/9781555819217.ch35.

Baril, T., Galbraith, J. and Hayward, A. (2024) ‘Earl Grey: A Fully Automated User-Friendly Transposable Element Annotation and Analysis Pipeline’, Molecular Biology and Evolution, 41(4), p. msae068. Available at: 10.1093/molbev/msae068.

Bendesky, A. et al. (2011) ‘Catecholamine receptor polymorphisms affect decision-making in C. elegans’, Nature, 472(7343), pp. 313–318. Available at: 10.1038/nature09821.

Berg, M. et al. (2016) ‘Assembly of the Caenorhabditis elegans gut microbiota from diverse soil microbial environments’, The ISME journal, 10(8), pp. 1998–2009. Available at: 10.1038/ismej.2015.253.

Blin, K. et al. (2021) ‘antiSMASH 6.0: improving cluster detection and comparison capabilities’, Nucleic Acids Research, 49(W1), pp. W29–W35. Available at: 10.1093/nar/gkab335.

Blum, M. et al. (2021) ‘The InterPro protein families and domains database: 20 years on’, Nucleic Acids Research, 49(D1), pp. D344–D354. Available at: 10.1093/nar/gkaa977.

Breger, J. et al. (2007) ‘Antifungal chemical compounds identified using a C. elegans pathogenicity assay’, PLoS pathogens, 3(2), p. e18. Available at: 10.1371/journal.ppat.0030018.

Cantalapiedra, C.P. et al. (2021) ‘eggNOG-mapper v2: Functional Annotation, Orthology Assignments, and Domain Prediction at the Metagenomic Scale’, Molecular Biology and Evolution, 38(12), pp. 5825–5829. Available at: 10.1093/molbev/msab293.

Červenák, F. et al. (2021) ‘Step-by-Step Evolution of Telomeres: Lessons from Yeasts’, Genome Biology and Evolution, 13(2), p. evaa268. Available at: 10.1093/gbe/evaa268.

Chan, P.P. et al. (2021) ‘tRNAscan-SE 2.0: improved detection and functional classification of transfer RNA genes’, Nucleic Acids Research, 49(16), pp. 9077–9096. Available at: 10.1093/nar/gkab688.

Conesa, A. and Götz, S. (2008) ‘Blast2GO: A Comprehensive Suite for Functional Analysis in Plant Genomics’, International Journal of Plant Genomics, 2008(1), p. 619832. Available at: 10.1155/2008/619832.

Cook, D.E. et al. (2017) ‘CeNDR, the Caenorhabditis elegans natural diversity resource’, Nucleic Acids Research, 45(D1), pp. D650–D657. Available at: 10.1093/nar/gkw893.

Crombie, T.A. et al. (2024) ‘CaeNDR, the Caenorhabditis Natural Diversity Resource’, Nucleic Acids Research, 52(D1), p. D850. Available at: 10.1093/nar/gkad887.

Davey, J.W. et al. (2020) Tapestry: validate and edit small eukaryotic genome assemblies with long reads, p. 2020.04.24.059402. Available at: 10.1101/2020.04.24.059402.

Deveau, A. et al. (2018) ‘Bacterial–fungal interactions: ecology, mechanisms and challenges’, FEMS Microbiology Reviews, 42(3), pp. 335–352. Available at: 10.1093/femsre/fuy008.

Dirksen, P. et al. (2016) ‘The native microbiome of the nematode Caenorhabditis elegans: gateway to a new host-microbiome model’, BMC Biology, 14(1), p. 38. Available at: 10.1186/s12915-016-0258-1.

Dirksen, P. et al. (2020) ‘CeMbio - The Caenorhabditis elegans Microbiome Resource’, G3 Genes|Genomes|Genetics, 10(9), pp. 3025–3039. Available at: 10.1534/g3.120.401309.

Emms, D.M. and Kelly, S. (2019) ‘OrthoFinder: phylogenetic orthology inference for comparative genomics’, Genome Biology, 20(1), p. 238. Available at: 10.1186/s13059-019-1832-y.

Falih, A.M. and Wainwright, M. (1995) ‘Nitrification, S-oxidation and P-solubilization by the soil yeast *Williopsis californica* and by *Saccharomyces cerevisiae*’, Mycological Research, 99(2), pp. 200–204. Available at: 10.1016/S0953-7562(09)80886-1.

Fang, J. et al. (2021) ‘Characteristics of a novel heterotrophic nitrification-aerobic denitrification yeast, *Barnettozyma californica* K1’, Bioresource Technology, 339, p. 125665. Available at: 10.1016/j.biortech.2021.125665.

Félix, M.-A. et al. (2011) ‘Natural and Experimental Infection of Caenorhabditis Nematodes by Novel Viruses Related to Nodaviruses’, PLOS Biology, 9(1), p. e1000586. Available at: 10.1371/journal.pbio.1000586.

Funannotate (2017). Available at: https://funannotate.readthedocs.io/en/stable/ (Accessed: 10 October 2024).

Gardes, M. and Bruns, T.D. (1993) ‘ITS primers with enhanced specificity for basidiomycetes- -application to the identification of mycorrhizae and rusts’, Molecular Ecology, 2(2), pp. 113–118. Available at: 10.1111/j.1365-294x.1993.tb00005.x.

Gonzalez, X. and Irazoqui, J.E. (2024) ‘Distinct members of the Caenorhabditis elegans CeMbio reference microbiota exert cryptic virulence that is masked by host defense’, Molecular Microbiology, 122(3), pp. 387–402. Available at: 10.1111/mmi.15258.

González-Hernández, R.A. et al. (2023) ‘Overview of fungal terpene synthases and their regulation’, World Journal of Microbiology & Biotechnology, 39(7), p. 194. Available at: 10.1007/s11274-023-03635-y.

Griem-Krey, H. et al. (2023) ‘The intricate triangular interaction between protective microbe, pathogen and host determines fitness of the metaorganism’, *Proceedings*. Biological Sciences, 290(2012), p. 20232193. Available at: 10.1098/rspb.2023.2193.

Groenewald, M. et al. (2023) ‘A genome-informed higher rank classification of the biotechnologically important fungal subphylum Saccharomycotina’, Studies in Mycology, 105(1), pp. 1–22. Available at: 10.3114/sim.2023.105.01.

Hasshoff, M. et al. (2007) ‘The role of Caenorhabditis elegans insulin-like signaling in the behavioral avoidance of pathogenic Bacillus thuringiensis’, The FASEB Journal, 21(8), pp. 1801–1812. Available at: 10.1096/fj.06-6551com.

Jiang, X., Xiang, M. and Liu, X. (2017) ‘Nematode-Trapping Fungi’, Microbiology Spectrum, 5(1). Available at: 10.1128/microbiolspec.FUNK-0022-2016.

Johnke, J. et al. (2025) ‘Caenorhabditis nematodes influence microbiome and metabolome characteristics of their natural apple substrates over time’, mSystems, 0(0), pp. e01533–24. Available at: 10.1128/msystems.01533-24.

Jones, J.D.G. and Dangl, J.L. (2006) ‘The plant immune system’, Nature, 444(7117), pp. 323–329. Available at: 10.1038/nature05286.

Kissoyan, K.A.B. et al. (2019) ‘Natural C. elegans Microbiota Protects against Infection via Production of a Cyclic Lipopeptide of the Viscosin Group’, Current Biology, 29(6), pp. 1030–1037.e5. Available at: 10.1016/j.cub.2019.01.050.

Kissoyan, K.A.B. et al. (2022) ‘Exploring Effects of C. elegans Protective Natural Microbiota on Host Physiology’, Frontiers in Cellular and Infection Microbiology, 12. Available at: 10.3389/fcimb.2022.775728.

Kobayashi, R., Kanti, A. and Kawasaki, H. (2017) ‘Three novel species of d-xylose-assimilating yeasts, Barnettozyma xylosiphila sp. nov., Barnettozyma xylosica sp. nov. and Wickerhamomyces xylosivorus f.a., sp. nov.’, International Journal of Systematic and Evolutionary Microbiology, 67(10), pp. 3971–3976. Available at: 10.1099/ijsem.0.002233.

Koren, S. et al. (2017) ‘Canu: scalable and accurate long-read assembly via adaptive k-mer weighting and repeat separation’, Genome Research, 27(5), pp. 722–736. Available at: 10.1101/gr.215087.116.

Kurtzman, C.P. (2011) The Yeasts: A Taxonomic Study. Elsevier.

Kurtzman, C.P., Robnett, C.J. and Basehoar-Powers, E. (2008) ‘Phylogenetic relationships among species of Pichia, Issatchenkia and Williopsis determined from multigene sequence analysis, and the proposal of Barnettozyma gen. nov., Lindnera gen. nov. and Wickerhamomyces gen. nov.’, FEMS Yeast Research, 8(6), pp. 939–954. Available at: 10.1111/j.1567-1364.2008.00419.x.

L, G. et al. (2024) ‘BRAKER3: Fully automated genome annotation using RNA-seq and protein evidence with GeneMark-ETP, AUGUSTUS and TSEBRA’, bioRxiv : the preprint server for biology [Preprint]. Available at: 10.1101/2023.06.10.544449.

Liu, K. et al. (2012) ‘How carnivorous fungi use three-celled constricting rings to trap nematodes’, Protein & Cell, 3(5), pp. 325–328. Available at: 10.1007/s13238-012-2031-8.

Lodder, J. (1932) ‘Über einige durch das “Centraalbureau voor Schimmelcultures” neuerworbene sporogene Hefearten’, Zentralbl. Bakteriol. Parasitenkd. Abt. II, 86, pp. 227– 253.

Lovell, J.T. et al. (2022) ‘GENESPACE tracks regions of interest and gene copy number variation across multiple genomes’, eLife. Edited by D. Weigel, 11, p. e78526. Available at: 10.7554/eLife.78526.

Macheleidt, J. et al. (2016) ‘Regulation and Role of Fungal Secondary Metabolites’, Annual Review of Genetics, 50(Volume 50, 2016), pp. 371–392. Available at: 10.1146/annurev-genet-120215-035203.

Madende, M. et al. (2020) ‘Caenorhabditis elegans as a model animal for investigating fungal pathogenesis’, Medical Microbiology and Immunology, 209(1), pp. 1–13. Available at: 10.1007/s00430-019-00635-4.

Meisel, J.D. and Kim, D.H. (2014) ‘Behavioral avoidance of pathogenic bacteria by *Caenorhabditis elegans*’, Trends in Immunology, 35(10), pp. 465–470. Available at: 10.1016/j.it.2014.08.008.

Mikheenko, A. et al. (2018) ‘Versatile genome assembly evaluation with QUAST-LG’, Bioinformatics, 34(13), pp. i142–i150. Available at: 10.1093/bioinformatics/bty266.

Minh, B.Q. et al. (2020) ‘IQ-TREE 2: New Models and Efficient Methods for Phylogenetic Inference in the Genomic Era’, Molecular Biology and Evolution, 37(5), pp. 1530–1534. Available at: 10.1093/molbev/msaa015.

Mullen, A.A. et al. (2018) ‘Draft Genome Sequences of Two Natural Isolates of the Yeast Barnettozyma californica from Ireland, UCD09 and UCD89’, Genome Announcements, 6(25), p. 10.1128/genomea.00548-18. Available at: https://doi.org/10.1128/genomea.00548-18.

Mylonakis, E. et al. (2002) ‘Killing of Caenorhabditis elegans by Cryptococcus neoformans as a model of yeast pathogenesis’, Proceedings of the National Academy of Sciences, 99(24), pp. 15675–15680. Available at: 10.1073/pnas.232568599.

Nakad, R. et al. (2016) ‘Contrasting invertebrate immune defense behaviors caused by a single gene, the Caenorhabditis elegans neuropeptide receptor gene npr-1’, BMC Genomics, 17(1), p. 280. Available at: 10.1186/s12864-016-2603-8.

Nordbring-Hertz, B., Jansson, H.-B. and Tunlid, A. (2011) ‘Nematophagous Fungi’, in *eLS*. John Wiley & Sons, Ltd. Available at: 10.1002/9780470015902.a0000374.pub3.

Nundaeng, S. et al. (2021) ‘An Updated Global Species Diversity and Phylogeny in the Genus Wickerhamomyces with Addition of Two New Species from Thailand’, Journal of Fungi, 7(11), p. 957. Available at: 10.3390/jof7110957.

Osman, G.A. et al. (2018) ‘Natural Infection of *C. elegans* by an Oomycete Reveals a New Pathogen-Specific Immune Response’, Current Biology, 28(4), pp. 640–648.e5. Available at: 10.1016/j.cub.2018.01.029.

Ou, S. et al. (2019) ‘Benchmarking transposable element annotation methods for creation of a streamlined, comprehensive pipeline’, Genome Biology, 20(1), p. 275. Available at: 10.1186/s13059-019-1905-y.

Pees, B. et al. (2024) ‘The Caenorhabditis elegans proteome response to two protective Pseudomonas symbionts’, mBio, 15(4), pp. e03463–23. Available at: 10.1128/mbio.03463-23.

Petersen, C. et al. (2023) ‘Host and microbiome jointly contribute to environmental adaptation’, The ISME Journal, 17(11), pp. 1953–1965. Available at: 10.1038/s41396-023-01507-9.

Pierce, E.C. et al. (2021) ‘Bacterial-fungal interactions revealed by genome-wide analysis of bacterial mutant fitness’, Nature microbiology, 6(1), pp. 87–102. Available at: 10.1038/s41564-020-00800-z.

Plissonneau, C., Stürchler, A. and Croll, D. (2016) ‘The Evolution of Orphan Regions in Genomes of a Fungal Pathogen of Wheat’, mBio, 7(5), p. 10.1128/mbio.01231-16. Available at: https://doi.org/10.1128/mbio.01231-16.

Pradel, E. et al. (2007) ‘Detection and avoidance of a natural product from the pathogenic bacterium Serratia marcescens by Caenorhabditis elegans’, Proceedings of the National Academy of Sciences, 104(7), pp. 2295–2300. Available at: 10.1073/pnas.0610281104.

Q, S. et al. (2023) ‘Large-Scale Detection of Telomeric Motif Sequences in Genomic Data Using TelFinder’, Microbiology spectrum, 11(2). Available at: 10.1128/spectrum.03928-22.

Reddy, K.C. et al. (2009) ‘A Polymorphism in npr-1 Is a Behavioral Determinant of Pathogen Susceptibility in C. elegans’, Science, 323(5912), pp. 382–384. Available at: 10.1126/science.1166527.

Samuel, B.S. et al. (2016) ‘Caenorhabditis elegans responses to bacteria from its natural habitats’, Proceedings of the National Academy of Sciences, 113(27), pp. E3941–E3949. Available at: 10.1073/pnas.1607183113.

Sato, T. et al. (2007) ‘The Rab8 GTPase regulates apical protein localization in intestinal cells’, Nature, 448(7151), pp. 366–369. Available at: 10.1038/nature05929.

Scherlach, K. and Hertweck, C. (2020) ‘Chemical Mediators at the Bacterial-Fungal Interface’, Annual Review of Microbiology, 74(1), pp. 267–290. Available at: 10.1146/annurev-micro-012420-081224.

Schulenburg, H. and Félix, M.-A. (2017) ‘The Natural Biotic Environment of Caenorhabditis elegans’, Genetics, 206(1), pp. 55–86. Available at: 10.1534/genetics.116.195511.

Schulenburg, H. and Müller, S. (2004) ‘Natural variation in the response of Caenorhabditis elegans towards Bacillus thuringiensis’, Parasitology, 128(Pt 4), pp. 433–443. Available at: 10.1017/s003118200300461x.

Schulte, R.D. et al. (2010) ‘Multiple reciprocal adaptations and rapid genetic change upon experimental coevolution of an animal host and its microbial parasite’, Proceedings of the National Academy of Sciences, 107(16), pp. 7359–7364. Available at: 10.1073/pnas.1003113107.

Shen, X.-X. et al. (2018) ‘Tempo and Mode of Genome Evolution in the Budding Yeast Subphylum’, Cell, 175(6), pp. 1533–1545.e20. Available at: 10.1016/j.cell.2018.10.023.

Shtonda, B.B. and Avery, L. (2006) ‘Dietary choice behavior in Caenorhabditis elegans’, Journal of Experimental Biology, 209(1), pp. 89–102. Available at: 10.1242/jeb.01955.

Simão, F.A. et al. (2015) ‘BUSCO: assessing genome assembly and annotation completeness with single-copy orthologs’, Bioinformatics, 31(19), pp. 3210–3212.

Snelders, N.C. et al. (2018) ‘Plant pathogen effector proteins as manipulators of host microbiomes?’, Molecular Plant Pathology, 19(2), pp. 257–259. Available at: 10.1111/mpp.12628.

Snelders, N.C. et al. (2020) ‘Microbiome manipulation by a soil-borne fungal plant pathogen using effector proteins’, Nature Plants, 6(11), pp. 1365–1374. Available at: 10.1038/s41477-020-00799-5.

Snelders, N.C. et al. (2021) ‘An ancient antimicrobial protein co-opted by a fungal plant pathogen for in planta mycobiome manipulation’, Proceedings of the National Academy of Sciences of the United States of America, 118(49), p. e2110968118. Available at: 10.1073/pnas.2110968118.

Sperschneider, J. and Dodds, P.N. (2022) ‘EffectorP 3.0: Prediction of Apoplastic and Cytoplasmic Effectors in Fungi and Oomycetes’, Molecular plant-microbe interactions: MPMI, 35(2), pp. 146–156. Available at: 10.1094/MPMI-08-21-0201-R.

Steenwyk, J.L. and Buida, T.J. (2022) Usage — biokit documentation. Available at: https://jlsteenwyk.com/BioKIT/usage/index.html#gene-wise-relative-synonymous-codon-usage-grscu.

Sun, J. et al. (2023) ‘OrthoVenn3: an integrated platform for exploring and visualizing orthologous data across genomes’, Nucleic Acids Research, 51(W1), pp. W397–W403. Available at: 10.1093/nar/gkad313.

Tan, M.-W., Mahajan-Miklos, S. and Ausubel, F.M. (1999) ‘Killing of Caenorhabditis elegans by Pseudomonas aeruginosa used to model mammalian bacterial pathogenesis’, Proceedings of the National Academy of Sciences, 96(2), pp. 715–720. Available at: 10.1073/pnas.96.2.715.

Tecle, E. and Troemel, E.R. (2022) ‘Insights from C. elegans into Microsporidia Biology and Host-Pathogen Relationships’, Experientia Supplementum (2012), 114, pp. 115–136. Available at: 10.1007/978-3-030-93306-7_5.

Tran, T.D. and Luallen, R.J. (2024) ‘An organismal understanding of *C. elegans* innate immune responses, from pathogen recognition to multigenerational resistance’, Seminars in Cell & Developmental Biology, 154, pp. 77–84. Available at: 10.1016/j.semcdb.2023.03.005.

Troemel, E.R. et al. (2008) ‘Microsporidia Are Natural Intracellular Parasites of the Nematode Caenorhabditis elegans’, PLOS Biology, 6(12), p. e309. Available at: 10.1371/journal.pbio.0060309.

Uliano-Silva, M. et al. (2023) ‘MitoHiFi: a python pipeline for mitochondrial genome assembly from PacBio high fidelity reads’, BMC Bioinformatics, 24(1), p. 288. Available at: 10.1186/s12859-023-05385-y.

White, T.J. et al. (1990) ‘38 - AMPLIFICATION AND DIRECT SEQUENCING OF FUNGAL RIBOSOMAL RNA GENES FOR PHYLOGENETICS’, in M.A. Innis et al. (eds) PCR Protocols. San Diego: Academic Press, pp. 315–322. Available at: 10.1016/B978-0-12-372180-8.50042-1.

Yang, Y. et al. (2007) ‘Evolution of nematode-trapping cells of predatory fungi of the Orbiliaceae based on evidence from rRNA-encoding DNA and multiprotein sequences’, Proceedings of the National Academy of Sciences, 104(20), pp. 8379–8384. Available at: 10.1073/pnas.0702770104.

Zárate-Potes, A., Schulenburg, H. and Dierking, K. (2024) ‘Unanticipated specificity in effector-triggered immunity’, Trends in Immunology, 45(12), pp. 939–942. Available at: 10.1016/j.it.2024.10.008.

Zhang, F. et al. (2017) ‘Caenorhabditis elegans as a Model for Microbiome Research’, Frontiers in Microbiology, 8. Available at: https://www.frontiersin.org/articles/10.3389/fmicb.2017.00485 (Accessed: 10 January 2024).

Zhang, F. et al. (2021) ‘Natural genetic variation drives microbiome selection in the Caenorhabditis elegans gut’, Current Biology, 31(12), pp. 2603–2618.e9. Available at: 10.1016/j.cub.2021.04.046.

Zhang, H. et al. (2018) ‘dbCAN2: a meta server for automated carbohydrate-active enzyme annotation’, Nucleic Acids Research, 46(W1), pp. W95–W101. Available at: 10.1093/nar/gky418.

Zhang, X. et al. (2021) ‘Antagonistic fungal enterotoxins intersect at multiple levels with host innate immune defences’, PLOS Genetics, 17(6), p. e1009600. Available at: 10.1371/journal.pgen.1009600.

Zimmermann, J. et al. (2024) ‘Gut-associated functions are favored during microbiome assembly across a major part of C. elegans life’, mBio, 15(5), p. e0001224. Available at: 10.1128/mbio.00012-24.

Zugasti, O. et al. (2016) ‘A quantitative genome-wide RNAi screen in C. elegans for antifungal innate immunity genes’, BMC Biology, 14(1), p. 35. Available at: 10.1186/s12915-016-0256-3.

Zugasti, O. and Ewbank, J.J. (2009) ‘Neuroimmune regulation of antimicrobial peptide expression by a noncanonical TGF-β signaling pathway in Caenorhabditis elegans epidermis’, Nature Immunology, 10(3), pp. 249–256. Available at: 10.1038/ni.1700.

